# Protein disorder controls allostery in DNA

**DOI:** 10.64898/2026.03.17.710445

**Authors:** Gabriel Rosenblum, Ivan Terterov, Sujeet Kumar Mishra, Nadav Elad, Tiberiu-Marius Gianga, Rohanah Hussain, Giuliano Siligardi, Hagen Hofmann

## Abstract

Metabolism, gene expression, and signaling all require the adaptation of protein activity to the mixture of reactants and products in a cell. This trait to adapt, called allostery, is hardwired in the structure of proteins. Binding a ligand at one location in a protein can change distant locations, thus tuning protein activity. How allostery works has been subject of intense research since its discovery sixty years ago. The challenge is to understand the order of events that follow ligand-binding in the three-dimensional architecture of proteins. Here we simplify this task by studying allostery in DNA, a nearly one-dimensional system. DNA can transmit allosteric signals over many nanometers to generate cooperativity in the binding of transcription factors, an archetype of the long-range action of allostery. We found that binding of the transcription factor ComK amplifies intrinsic microsecond structural fluctuations in DNA many nanometers distant from the binding site. Yet, it is not protein binding per se, but the intrinsically disordered region (IDR) of the protein that amplifies these fluctuations. IDR removal does not only rigidify DNA, but it also abolishes allostery. The result is a structurally distorted protein-DNA complex that lost its function. These findings have important implications for our understanding of transcription activation and suggest a new functional role for IDRs in transcription factors.

---

Few ideas in biology are general enough to apply to all life on earth. The idea of allostery is one of them. Allostery is the coupling between two distant sites in a biomolecule^1-3^. Binding a ligand to one site changes the remote site and tunes its function. Remote sites can bind other ligands, or they are active centers in enzymes. Without this property, life would cease to exist. While the thermodynamic prerequisites of allostery in proteins have been worked out^4^, it has been incredibly challenging to study how exactly allosteric signals travel nanometer distances in biomolecules due to the intertwined topology of structured proteins. However, allostery is also a property of DNA, which is a rather recent discovery^5-21^. The simplicity of linear double stranded DNA (dsDNA) promises an easier assessment of the question. At short length scales, less than hundred base pairs (bp), dsDNA is stiff, leaving only two directions in which a signal can possibly ‘travel’. The price of this simplicity is that structural changes are limited to moderate bending or twisting of the double helix. But also structural changes in proteins are often small, so small that they even fall short in explaining allostery^22-24^. These cases triggered a revival of the idea of ‘dynamic allostery’^25,26.^ Different from classical models^27-29^, the first ligand would change the thermal fluctuations of a biomolecule rather than its average structure, which then allows a second ligand to bind with greater affinity^4^. Clear-cut demonstrations of dynamic allostery are scarce because of the experimental challenges associated with measuring structural fluctuations in proteins. Catabolite activator protein (CAP)^30,31^, thymidylate synthase^32^, and the glucagon receptor^33^ are rare examples. For DNA, direct experimental evidence for dynamic allostery has so far been missing altogether. Two cases in which allostery in DNA has been demonstrated are known^5,19,^ but only one of them is found in nature^19^: The bacterium *Bacillus subtilis* requires an all-or-none binding of the transcription factor (TF) ComK to its promoter for the stochastic switch into a competent phenotype in which the bacteria can import DNA^34-36^. The promoter is inactive at low ComK concentrations but switches cooperatively to an active state within a narrow concentration range^34,36^ (Fig. 1a). We showed previously that this switch is cooperative because binding of ComK to one location increases its affinity to another site many nanometers away^19^. This raises the fundamental question of how exactly the information of binding to the first site has made its way to the second site.

**Figure 1.**
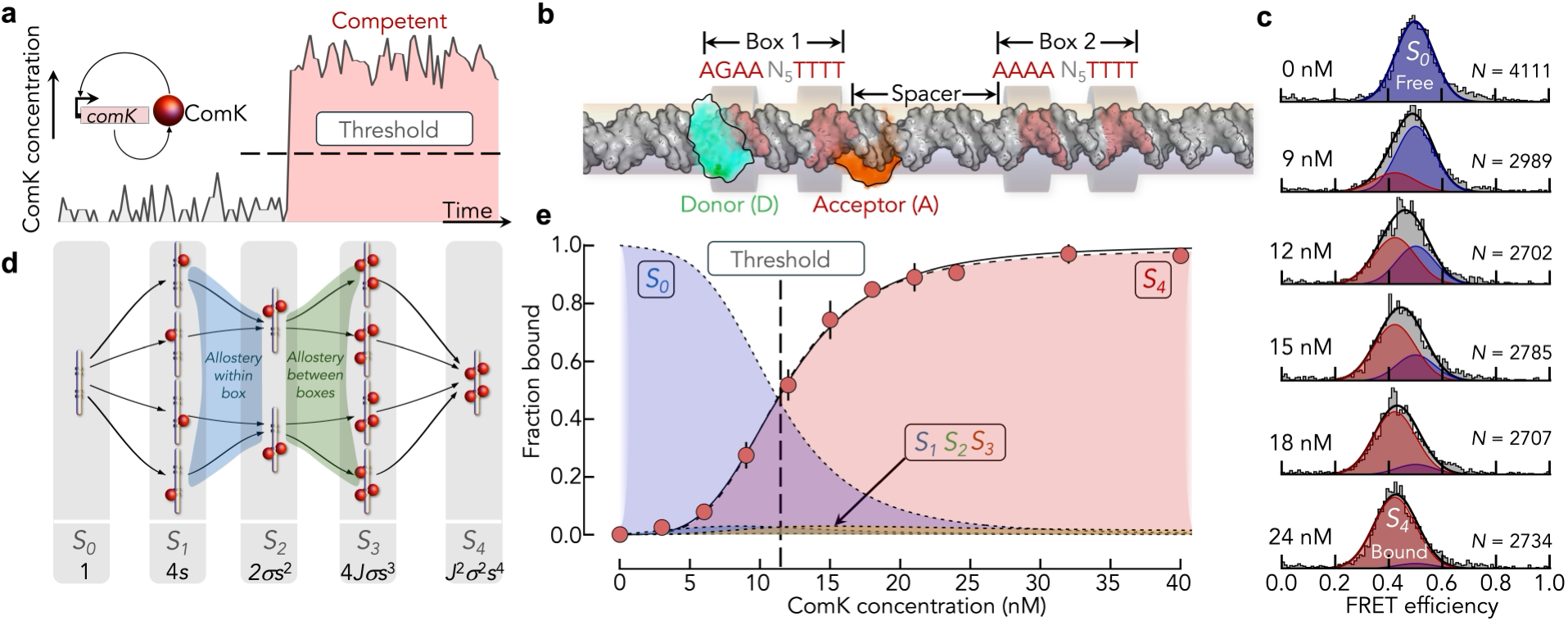
Allostery in the ComK promoter. **(a)** When fluctuations in the concentration of ComK cross a certain threshold in *B. subtilis*, ComK binds stably to its promoter and initiates the expression of the competence genes. Inset: Positive feedback loop in the gene expression mechanism of ComK. **(b)** The promoter has two boxes linked by a spacer. Each box binds two ComK molecules. The donor (D, green) and acceptor (A, orange) dyes are attached to probe the distance in box 1. **(c)** FRET efficiency histograms of DNA molecules at increasing concentrations of ComK (indicated). Solid lines are fits with two Gaussian peaks that represent free DNA (S_0_) and DNA bound with four ComKs (S_4_). The number of molecules (*N*) is indicated. **(d)** Allosteric Pauling-model of the binding of ComK (red ball) to its promoter (gray tube). The statistical weights of the five macro-states (*S*_*0*_…*S*_*4*_) are given at the bottom. Here, *s* = *K* [ComK] where *K* is the association constant of a single monomer to any of the four binding sites. The parameters *σ* and *J* describe the cooperativity within and between boxes, respectively. Micro-states within each macro-state are schematically shown within the gray bars. Based on previous cryo-EM results^19^, *S*_*2*_ only consists of two micro-states. **(e)** Average fraction of ComK-bound DNA molecules as function of the ComK concentration (circles) from 2 independent measurements. The tips of the black vertical lines indicate the values of 2 independent repeats. Solid line is a fit with the Hill function. The fit (dashed line) with a Pauling binding model is shown together with the predicted fraction of the macro-states of the model (dotted lines).

To answer this question, we combined single-molecule Förster resonance energy transfer (smFRET) and cryogenic electron microscopy (cryo-EM) with coarse-grained (CG) molecular simulations and elastic network models (ENMs). We found that binding of ComK to the first location amplifies thermal microsecond structural fluctuations of DNA. This flexibility decays with the distance to the binding site. We explain this flexibility by weak interactions of an intrinsically disordered region (IDR) of ComK with the DNA. Recently, the role of IDRs in TFs has gained much attention^37^. While previously thought to be responsible for recruiting auxiliary factors in transcription, evidence mounts that IDRs interact directly with DNA, thus impacting and potentially even modulating motif search across genomes and DNA binding affinities^38^. Our results show that IDRs can control allosteric signal transmission via DNA, which is a previously unknown function of IDRs.

The distant recognition sites for ComK on DNA are termed box 1 and box 2 (Fig. 1b). Each box binds two ComK molecules^19^. The boxes are separated by spacers of variable length and only three spacers of 8, 18, and 31 base pairs (bp) are found in *Bacillus subtilis*^39^. The sequence within a box contains two A-tracts, which are stretches of four consecutive adenine bases that bend the DNA towards the minor groove^19,40^. To demonstrate the allosteric effect in this system, we chose a promoter with an 18bp spacer and attached a donor and an acceptor fluorophore at the beginning and at the end of box 1 to probe the structural changes in freely diffusing DNA with smFRET (Fig. 1b, Extended Data Table 1). Addition of ComK, which is monomeric under our conditions^19^ (Extended Data Fig. 1), shifts the distribution of FRET efficiencies to lower values, which translates into a ∼3Å increase in the distance between donor and acceptor (Fig. 1c). We fitted the FRET efficiency distributions to a sum of two Gaussian peaks to derive the fraction of ComK-DNA complexes (Fig. 1c, Methods). This fraction (*f*) increases in a sigmoidal manner with the concentration (*x*) of ComK^19^. With *K*_*D*_ being the apparent dissociation constant of a single ComK molecule, a fit with the Hill equation 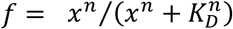 results in a Hill exponent of *n* = 3.7 ± 0.2, i.e. we probe the binding of at least four ComK molecules. The same experiment only gave *n* = 1.7 ± 0.1 for a promoter with a single box (Extended Data Fig. 2a,c), in accord with two ComKs per box. Since we found four ComKs in the two-box promoter even though we only monitored one of the two boxes, the boxes must communicate allosterically. A mechanistic description of the binding reaction with a Pauling allostery model^41^ (Fig. 1d,e) allowed us to determine the free energy of coupling between both boxes, which amounts to Δ*G*_*J*_ = −5.6 ± 1 *k*_*B*_*T* (Methods, Extended Data Table 2), where *k*_*B*_ is the Boltzmann constant and *T* is the room temperature (296 K). Hence, the affinity of ComK for a box is 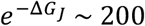 fold higher if the other box is already bound by two ComKs, the archetype of allostery.

Early *in vivo* experiments with *B. subtilis* showed that the removal of 35 amino acids from the C-terminus of ComK abolishes the transcriptional activity^42^. This is surprising as this tail is predicted to be an intrinsically disordered region (IDR). In fact, not only has the helical signature of the tail in the AlphaFold2 prediction a low confidence, but it is invisible in our previously published cryo-EM map of the ComK-DNA complex (Fig. 2a), indicating that it is highly flexible^19^. When we used synchrotron radiation circular dichroism (SRCD, Methods), we find that the isolated IDR is indeed largely unstructured with only 18% helical signature (Fig. 2b). However, when we measured the binding cooperativity of ComK without this IDR (ComKΔ) to a two-box promoter and compared it with the wildtype (Fig. 2c), we found a strongly reduced Hill exponent of *n* = 1.8 ± 0.1 (Fig. 2d), which is identical to that found for the one-box promoter with either ComKΔ (*n* = 1.6 ± 0.1) or wildtype ComK (*n* = 1.7 ± 0.1) (Extended Data Fig. 2a-d). Since ComKΔ is also monomeric under our experimental conditions (Extended Data Fig. 1), this value means that the cooperativity within the dimer in a box is unaffected by the IDR-removal while the cooperativity between boxes is lost. Expectedly, a fit with the Pauling model results in a zero free energy of coupling between the boxes (Fig. 2e). This loss of cooperativity is independent of the spacer length. While the Hill exponent and the coupling free energy oscillate with the periodicity of the DNA double helix for ComK, the removal of the IDR abolished cooperativity at all spacer lengths (Fig. 2f,g). How can we explain the breakdown of cooperativity in ComKΔ?

**Figure 2.**
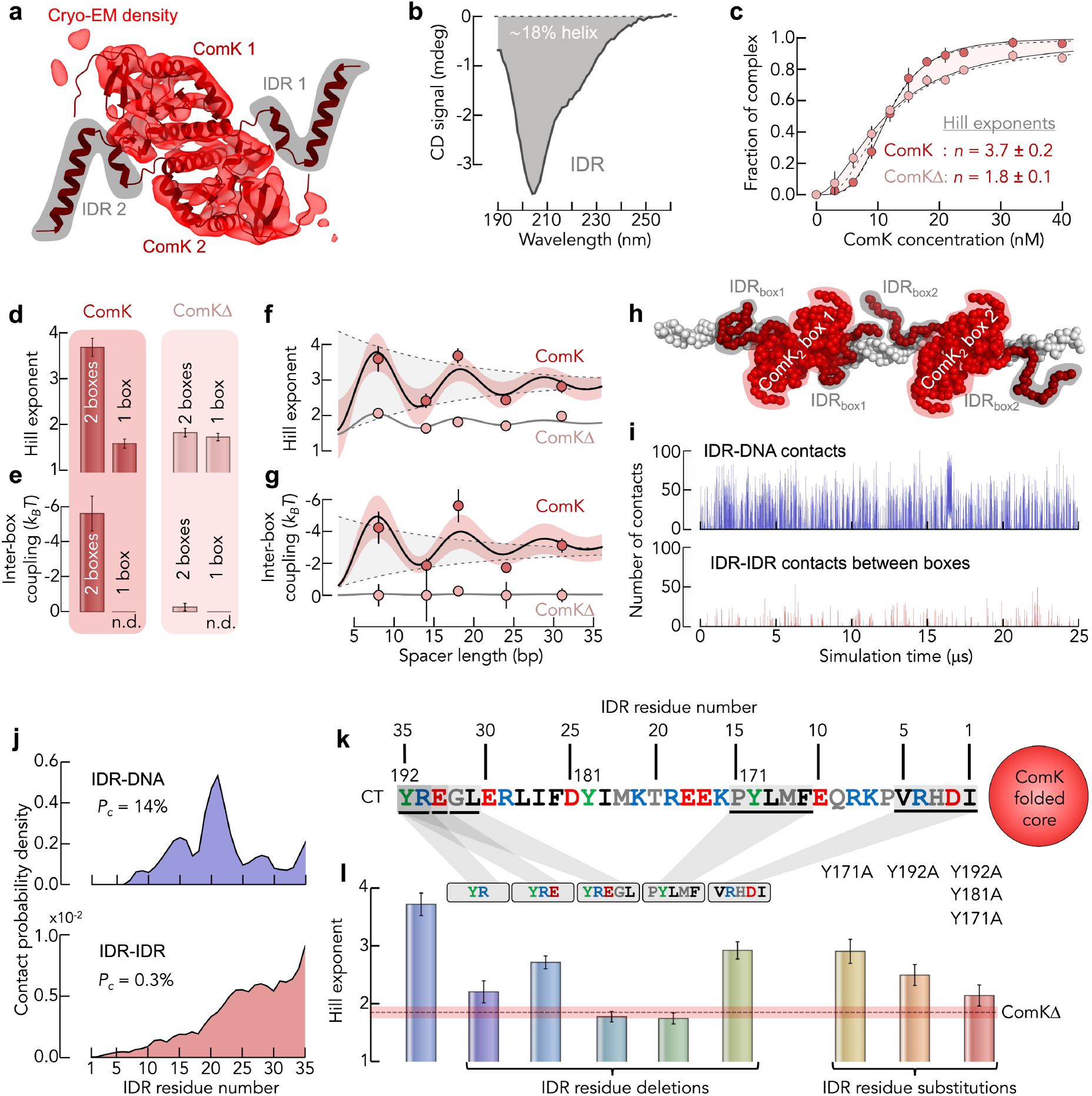
The IDR of ComK modulates allostery. **(a)** Cryo-EM density of a ComK dimer (light red) bound to a box in the DNA (not shown) in comparison to the AlphaFold2 prediction (dark red). The gray shaded area highlights the IDR of ComK. **(b)** Synchrotron radiation CD spectra of the isolated IDR of ComK. **(c)** Comparison of the binding isotherm of ComK (dark red circles) and that of a variant without the IDR (ComKΔ) (light red circles). The tips of the black vertical lines indicate the values of 2 independent repeats. Solid lines are fits with the Hill function and dashed lines are fits with the Pauling model. The Hill exponents are given as inset. **(d, e)** Hill exponents (d) and coupling free energies (e) for promoters with two and one box for ComK and ComKΔ. Colors are indicated. Error bars are the standard error from a global fit of two independent data sets. **(f, g)** Hill exponents (f) and coupling free energies (g) for ComK and ComKΔ as function of the spacer lengths. Solid lines are empirical fits of an exponentially damped oscillation with a decay length of 14bp determined in ref.^19^. Error bars are the standard error from a global fit of two independent data sets. **(h)** CG-model of the ComK-DNA complex. **(i)** Trajectory of number of contacts between IDRs and DNA (top) and between the IDRs of distal boxes (bottom) from the CG-simulations. **(j)** Contact probability distribution of contacts between IDRs and DNA (top) and of contacts between distal IDRs (bottom). The overall contact probability is indicated. **(k)** Amino acid sequence of the C-terminal IDR of ComK. **(l)** Hill exponents of ComK variants with modified IDRs (underscore indicates deletion). Vertical lines indicate the error from global fits to two independent experiments for each variant.

In the simplest model, allostery would arise from contacts between IDRs from opposite boxes. Allostery would merely result from a ‘bridge’ formed by the IDRs between the boxes. However, molecular dynamics simulations using a coarse-grained (CG) model of the ComK-DNA complex challenge this conclusion. In the simulations, we kept the structure of DNA and the folded core of ComK static (Methods) to estimate the probability that IDRs from opposite boxes contact each other (Fig. 2h,i). We found that the IDRs from opposite boxes reach each other only rarely with a probability <0.3% (Fig. 2j). The formation of an IDR-IDR “bridge” is therefore unlikely. This is in line with other findings, for instance with the fact that even a long 31bp spacer still shows significant cooperativity (Fig. 2f), or our previously published result that a single-strand break of the DNA in the spacer also abolishes allostery^19^, which cannot be explained with IDR-IDR contacts and rather points to a key role for DNA in the allosteric mechanism. When we truncate the IDR partially (Extended Data Table 2), we obtained an unexpected result (Fig. 2k). The removal of the terminal two and three amino acids reduces allostery and deleting the terminal five amino acids or five amino acids in the central region of the IDR abolishes inter-box allostery entirely (*n* = 1.7 ± 0.1) (Fig. 2l). However, it is not the reduced length of the IDR per se that eliminates allostery. Another truncation of five amino acids at the N-terminus of the IDR still shows cooperativity (*n* = 2.9 ± 0.2) (Fig. 2l), albeit to a reduced extent. Moreover, when we replaced three tyrosine residues of the IDR by alanine without truncating the tail, the allostery was also substantially reduced (*n* = 2.1 ± 0.2). Hence, IDR sequence rather than length seems the decisive parameter. What else do the IDRs contact? Our coarse-grained simulations indicate a plethora of contacts of the IDR with DNA (Fig. 2i,j), raising the question of whether the IDRs impact DNA properties?

We therefore determined the cryo-EM structure of ComKΔ in complex with the 18bp spacer promoter and compared it to our previously published structure of the ComK-DNA complex^19^ (Methods). We found three particle classes: DNA, DNA with a ComKΔ-dimer at one box and DNA with a ComKΔ-dimer at each box (Fig. 3a). The relative abundance of these classes allows us to calculate the Hill exponent (*n*_*EM*_) directly from the cryo-EM experiment (Methods). We found *n*_*EM*_ = 2.3 with ComKΔ compared to *n*_*EM*_ = 4.0 with wildtype ComK^19^, i.e. in line with the reduced cooperativity found with smFRET. The lack of particles with just a single ComKΔ per box also indicates that the cooperativity within a box is retained, in line with the results from our smFRET experiments. To identify structural changes upon IDR-removal, we collected the particles with both boxes bound by ComKΔ (73,689) and determined a 3D cryo-EM map at a resolution of 6Å. The electron density of ComKΔ in each box is very similar to that obtained previously with wildtype ComK (Fig. 3b). In fact, the AlphaFold2 structure of wildtype ComK fits the cryo-EM maps of both ComK and ComKΔ perfectly with only marginal differences at the length-scale of a single box. However, at the length scale of the full promoter (93bp), the DNA topology differs strongly between the complexes (Fig. 3c). The DNA is straight with only moderate bending with wildtype ComK. With ComKΔ on the other hand, the promoter is twisted and bent. Apparently, the IDR reshapes the DNA structure at nanometer length scales. Even more so, a 3D variability analysis^43^ (Methods) shows a higher variability in the DNA-ComK complex compared to the DNA-ComKΔ complex (Fig. 3d), indicating that the IDR does not only alter the DNA structure but also its flexibility. Whether the straight DNA topology in presence of the IDR is origin or result of the increased flexibility is unclear at this point. What is clear though is that our inferences about flexibility rely on experiments in which the samples were quenched to cryogenic conditions of ∼90K. If the timescales associated to this flexibility are faster than the cooling to cryogenic temperatures, conformations will partially equilibrate^44^, thus providing 3D-maps that do not reproduce the ensemble at physiological temperatures. We therefore measured the flexibility of DNA, i.e. amplitudes and timescales of structural fluctuations, directly at room temperature.

**Figure 3.**
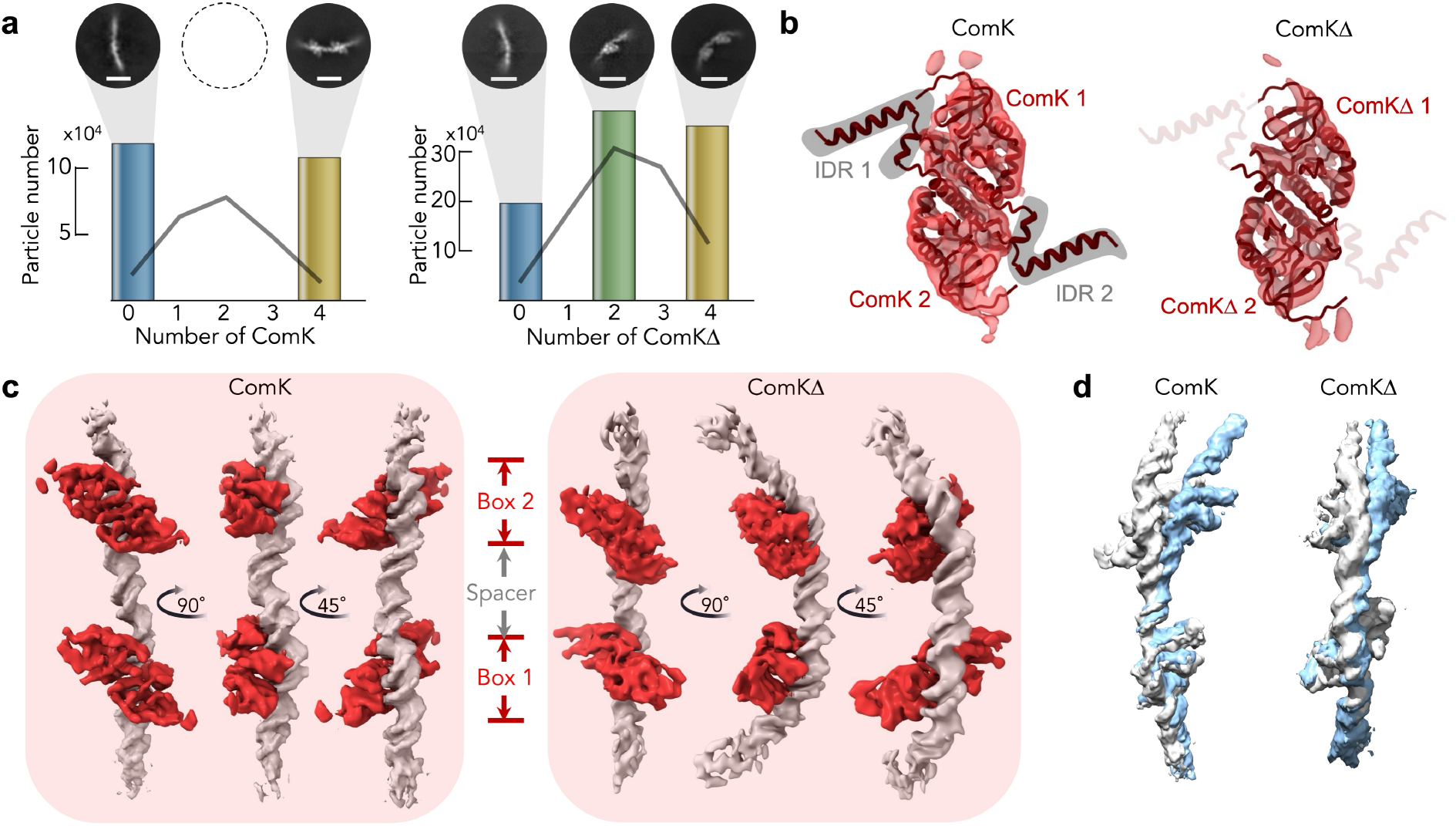
Comparison of the cryo-EM densities of ComK and ComKΔ. **(a)** Distribution of protein-DNA complexes with different numbers of ComK (left) and ComKΔ (right) bound for a promoter with an 18 bp spacer. The class averages of the particle types are shown on top. **(b)** Comparison of the cryo-EM densities of ComK (left) and ComKΔ (right) and best fit of the AlphaFold2 model of full-length ComK including the IDR. The IDR is indicated in gray. **(c)** Cryo-EM density of the ComK-DNA complex (left, EMD-11022) and the cryo-EM density of the ComKΔ-DNA complex (right, EMD-56533). **(d)** Result of the 3D-variability analysis using CryoSPARC of the ComK-DNA complex (left) and the ComKΔ-DNA complex (right). The two extremes of the variability components were overlaid to demonstrate the reduced variability in ComKΔ-DNA. The angle of view was chosen to maximize the visual difference between the two extreme components in both complexes.

We first determined the magnitude of fluctuations in the distance between our FRET donor and acceptor dyes using the shortest timescale available in smFRET experiments: the nanosecond fluorescence lifetime of our fluorophores (∼4ns). If much slower than the fluorescence lifetime, motions in the DNA would appear as a ‘frozen’ distribution of donor-acceptor distances in these experiments (Fig. 4a-d). The width of this distribution measures the amplitude of structural fluctuations. We labelled the two-box promoter with donor and acceptor in three regions: box1, spacer (18bp), and box2 and recorded the fluorescence lifetime decays of the donor and the acceptor (Methods, Extended Data Fig. 4 and 5). In each experiment, we selected molecules with active donor and acceptor dyes (Fig. 4e). For a single static distance, the donor decay would be exponential (Fig. 4d,f). However, we found a highly non-exponential decay, indicative of a distribution of distances (Fig. 4f) together with an initial rise in the decay of the acceptor (Fig. 4g). Using a distance distribution model with position and width parameter (Methods), we described the donor and acceptor decays in a global manner and obtained excellent fits (Fig. 4f,g, Extended Data Table 4). We then determined the width of the distance distribution in excess to the variability due to the aliphatic linkers of our dyes, which we previously estimated with a rotational isomeric state model^19^. In free DNA, the excess distance fluctuations have an amplitude of 1.8±0.6Å when averaged over the three probed regions (Fig. 4h). In the boxes, binding of either ComK or ComKΔ increases these fluctuations to 5.2±0.3Å on average. So, both boxes are more flexible in the presence of either variant. However, in the spacer between the boxes, the fluctuations with wildtype ComK (8.4±0.1Å) are clearly stronger than with ComKΔ (6.0±0.1Å) (Fig. 4i). It therefore is the spacer between the boxes whose flexibility is most affected by the IDR. A disadvantage of our determination of the fluctuation amplitude is its dependence on the chosen model of the distance distribution. To obtain an independent and model-free determination of the fluctuation amplitude together with the timescale of these fluctuations, we computed the correlation function of FRET efficiencies using our recently developed method, termed FECS (FRET Efficiency Correlation Spectroscopy)^45^ (Methods, Extended Data Table 5). The FECS-functions decay over microsecond timescales with two characteristic timescales, 5±1μs and 154±15μs (Fig. 4j, Extended Data Fig. 6). Although the faster timescale is close to the expected lifetime of the photophysical triplet states of our FRET dyes, its amplitude exceeds the expected triplet amplitude by far^45^, suggesting substantial DNA motions even at this fast timescale. The amplitudes of the FECS-functions reproduce the trend of the distance distribution widths from the fluorescence lifetimes: fluctuation amplitudes are similar for ComK and ComKΔ in the boxes, but in the spacer, they are much enhanced in the presence of the IDR (Fig. 4j, middle). A dynamic allostery model^26,46^ can now be pictured in which ComK-binding to the first box increases the flexibility of the DNA, which loosens steric constraints at the distant box and permits a preferential binding of the second pair of ComKs. However, this idea requires that already the binding of two ComKs to a single box increases DNA flexibility. We therefore performed smFRET experiments with a one-box promoter in which we probe the flexibility of DNA at increasing distances from box (Fig. 5a,b). Both, the distance distributions obtained from the fluorescence lifetime experiments and the amplitudes of the FRET correlation functions (Fig. 5c) show enhanced flexibility of the DNA in the presence of wildtype ComK. With ComKΔ, this flexibility is reduced. Most importantly, the flexibility decays with increasing distance from the binding site (Fig. 5b). Using an exponential decay, we estimated the characteristic lengths scale of this decay to be 15±4 bp, which coincides with the length scale of allosteric signal transmission of 14±8 bp (Fig. 2f).^5,19^,47 But how does the IDR control DNA flexibility if persistent contacts between IDR and DNA are clearly missing in the cryo-EM map? Our simulations suggest that transient interactions with the DNA are responsible for it. To test this possibility, we measured the contact-formation between IDR and DNA using quenching between our donor dye (AlexaFluor488) and a ComK variant with a tryptophan residue at the terminus of the IDR (Y192W) (Fig. 5d, Methods). It has previously been shown that tryptophan quenches AlexaFluor488 via the formation of a stacked complex (static quenching)^48^. If the fluorescence intensity of the dye is unaffected by the presence of the IDR, we can exclude the possibility of IDR-DNA contacts as origin of the increased DNA-flexibility. We placed the dye at different locations on the DNA and recorded the fluorescence of the dye. When we compared the fluorescence signals between wildtype ComK and ComKΔ, we found that quenching was indeed observed in presence of the IDR across the entire probed length scale of 18bp in both directions 5’-3’ and 3’-5’ (Fig. 5e). The magnitude of quenching was very small with only 10% reduced fluorescence in the presence of the IDR. Despite frequent interactions of the IDR with the DNA, the probability that the specific C-terminal tryptophan contacts a specific location on the DNA is also very low in the simulations. However, when rescaled, the simulated quenching profile qualitatively matches the experimental data, indicating IDR and DNA do come in proximity (Methods). Hence, interactions between IDR and DNA therefore seem to be a possible explanation for the enhanced flexibility of DNA. We therefore postulate that transient contacts between IDR and DNA destabilize the double helix locally thus mediating allostery between both binding boxes.

**Figure 4.**
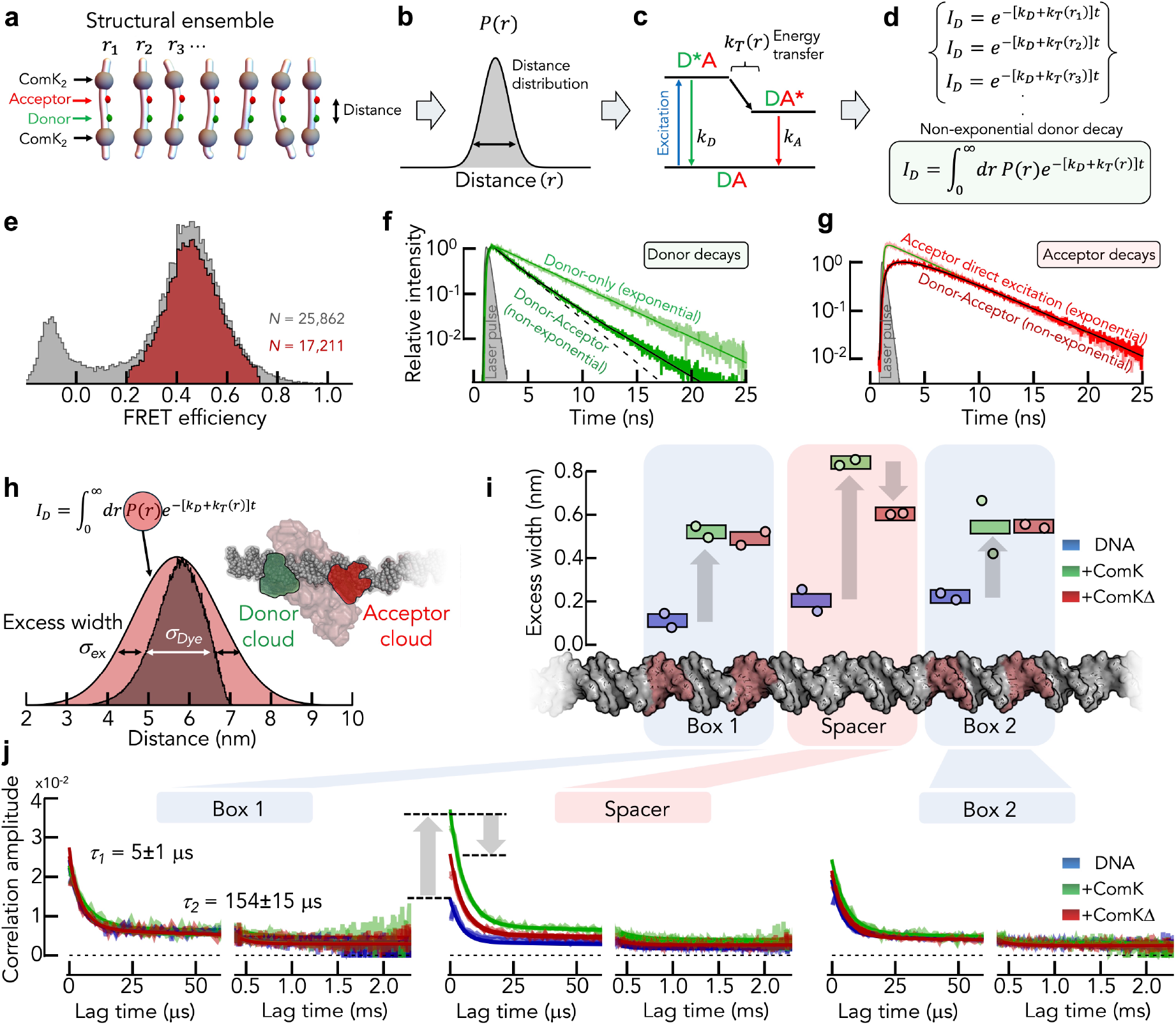
Determining amplitudes and timescales of structural fluctuations in DNA. **(a-d)** Schematic depiction of how a distance distribution impacts the fluorescence lifetime decay of the donor dye. A structural ensemble of ComK-promoters (here labeled in the spacer) (a) leads to a distribution of distances (b). The photophysical cycle in FRET depends on the kinetic rate constants of emission (*k*_D_, *k*_*A*_) and on the energy transfer rate *k*_*T*_ that is a function of the distance *r* between the dyes (c). This leads to a slightly different lifetime decay for each distance in the ensemble (d). The weighted sum (integral) of all these decays is the experimentally observed lifetime decay of the donor. **(e)** smFRET histogram of the 18bp spacer promoter in the presence of 100 nM ComK before (gray) and after (red) the elimination of molecules with bleached acceptor (Methods). **(f)** Fluorescence lifetime decay of the donor in the presence (dark green) and in the absence (light green) of the acceptor. Solid lines are fits with the distance distribution model (dark green) or an exponential decay (light green). The dashed black line indicates a fit of an exponential decay if only the first 4 ns of the donor decay were included in the fit. **(g)** Fluorescence lifetime decay of the acceptor after donor excitation (dark red) and after direct excitation (light red). Solid lines are fits with the distance distribution model (dark red) or an exponential decay (light red). **(h)** Comparison of the fitted distance distribution from f and g compared to the distribution of distance that results from the flexible aliphatic linkers of our dyes. Inset: Schematic that illustrates the extend of the positional distribution of the FRET dyes due to the dye-linker flexibility. **(i)** Excess width of distance fluctuations in the DNA for three regions: box1, spacer, box2 determined from the fluorescence lifetime decays. Rectangles indicate the mean of two independent experiments (circles). Colors are indicated. **(j)** Determination of the timescale of distance fluctuations using FRET efficiency correlation functions (FECS) in the three regions: box1, spacer, box2. Solid lines are fits with two exponential decays and an offset that accounts for static heterogeneity. The average of the decay times over all experiments are indicated.

**Figure 5.**
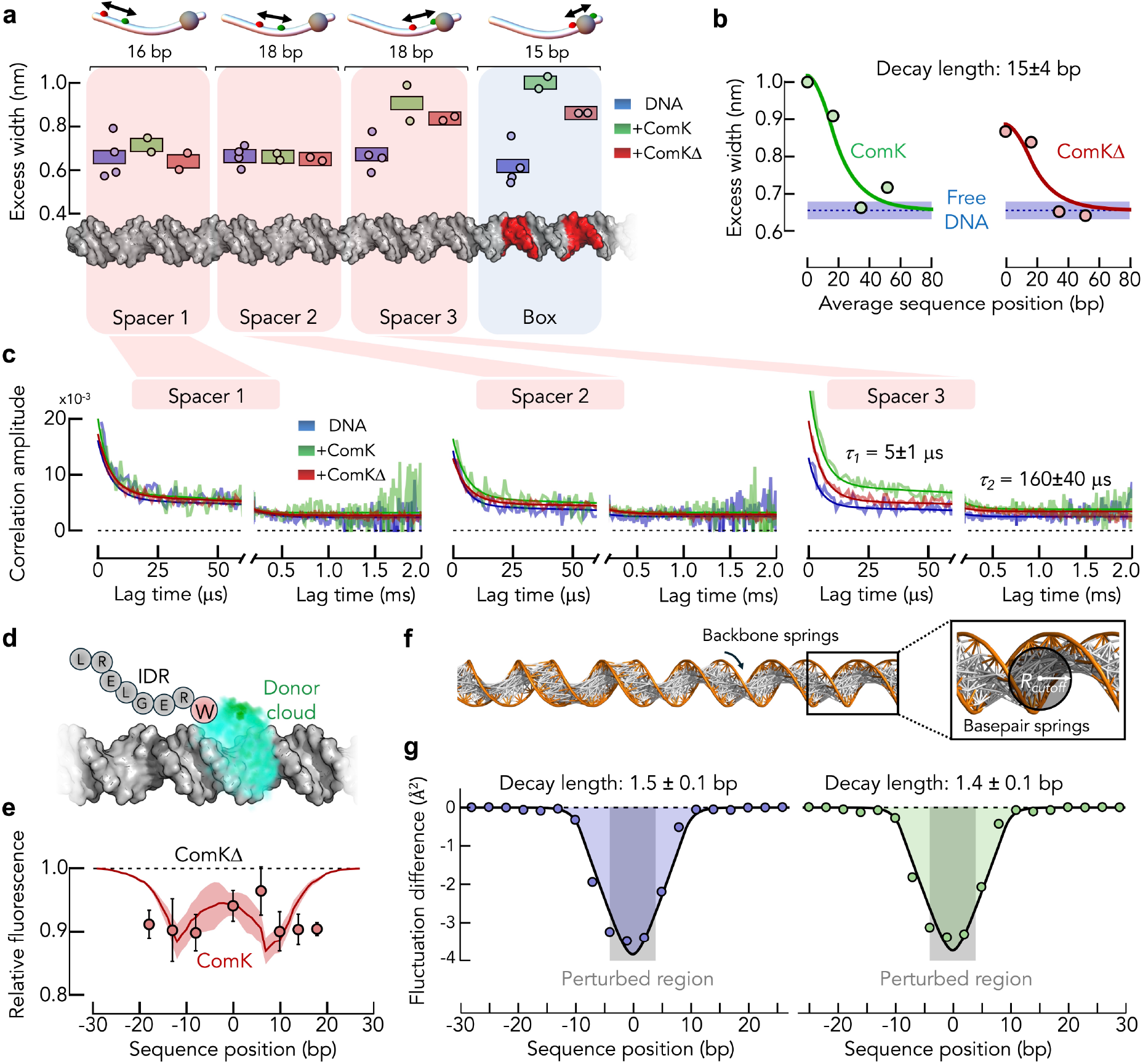
Propagation of fluctuations along the DNA. **(a)** Excess width of distance fluctuations in the DNA for a promoter with one box, determined from the fluorescence lifetime decays. Rectangles indicate the mean of at least two independent experiments (circles). Colors are indicated. **(b)** Decay of the excess-width as function of the distance from the box for ComK and ComKΔ. Solid lines are global fits with an exponential decay, convoluted with a box function to account for the averaging across the probed distances. The global decay length is indicated. **(c)** Determination of the timescale of distance fluctuations using FECS at increasing distance from the box. Solid lines are fits with two exponential decays and an offset that accounts for static heterogeneity. The average of the decay times over all experiments are indicated. **(d)** Scheme of the static quenching of the donor AlexaFluor488 attached to the DNA (shown as a positional cloud due to the flexibility of the dye linker) by an engineered tryptophan (Y192W) at the C-terminus of the IDR. **(e)** Relative bulk fluorescence intensities of a ComK-DNA complex for a promoter with one box relative to the intensities obtained for the ComKΔ-DNA complex (circles). The sequence position of the dye-location is given relative to the center of the box. Error bars are ±1SD with >3 independent experiments. Solid line is the re-scaled probability of the dye not being in contact with tryptophan (scaling factor 52) from the CG-simulation. The shaded band indicates ±2SD over all simulation runs. **(f)** Representation of B-DNA in the elastic network model. Each connection represents a harmonic spring. Springs connecting the backbone are shown in orange. Springs representing the base pairs (gray) were placed by connecting all DNA beads within a cutoff radius of *R*_*cutoff*_ = 10 nm. **(g)** Change in the distance fluctuations across 9bp in a 1000bp long DNA after increasing the spring constant fivefold (blue) and threefold (green) in a central 9bp-region (gray bar). The solid line (black) is a fit with an exponential decay convoluted with a box-function of 9bp in width. The decay length is indicated.

In fact, a recent model of DNA-mediated allostery supports the idea of an increase in DNA-flexibility by ComK^20^. How far can local perturbations possibly propagate along regular B-DNA? A bead-and-spring representation of B-DNA in form of an elastic network model (ENM, Methods)^49-51^ provides a rough estimate. Whereas the ComK-promoter has specific sequence features and a bend topology, we are more interested in understanding whether B-DNA is generally capable of propagating local changes in flexibility purely based on its intrinsic elasticity. We used a long DNA (1000 bp) to minimize potential finite-length effects. The DNA-model consisted of three beads for each nucleotide (base, sugar, and phosphate) that were connected using harmonic springs (Fig. 5f). To produce a local perturbation, we changed the spring stiffness in a local region of 9bp, which represents a perturbed box. Using normal-mode decomposition, we then determined the expected distance-fluctuation amplitude distant from the perturbed region. We found a decay length scale of only 1.5 bp along the double helix (Fig. 5g). This value is at odds with the value found experimentally (15±4 bp), indicating that the IDR of ComK generates flexibility that exceeds the elastic properties expected for the ideal structure of B-DNA. This excess flexibility might originate from partial dissociations of base pairs and/or sequence-specific features such as the intrinsic bending-properties of the A-tracts in the promoter of ComK. In summary, our results demonstrate that allostery in the binding of ComK to its promoter requires the IDR of the protein. Correlated with the removal of the IDR are (i) a distorted structure of the protein-DNA complex and (ii) reduced microsecond structural fluctuations of the DNA. Given the prevalence of IDRs in transcription factors, dynamic allostery caused by weak and transient contacts of IDRs with DNA might be a general mechanism to generate binding cooperativity in gene expression processes.

## Discussion

Most eukaryotic transcription factors contain long stretches of IDRs that can reach hundreds of residues in length^52-59^. These disordered sequences seem to evolve rapidly, even if they are functionally conserved^60-66^. Nevertheless, recent studies demonstrated that such IDRs are strictly required for the specificity of dozens of TFs in budding yeast^67-69^. Yet, how IDRs confer this specificity is unclear. Possible mechanisms range from triggering phase separation of the transcription machinery at specific genomic locations^70,71^ over mediating interactions with cofactors or chromatin, up to autoinhibitory functions via intramolecular contacts with the folded DNA-binding domain. We found that a comparatively short IDR of only 35 amino acids has a pronounced impact on the structure and dynamics of DNA. Especially the flexibility of DNA is increased at nanometer lengths scales by this IDR. We showed that this flexibility is strongly correlated with the allosteric coupling between two distant binding sites that allows the bacterial transcription factor ComK to bind with high cooperativity to its promoter. Key for this destabilization of DNA seem to be direct contacts between IDR and DNA whereby aromatic residues such as tyrosine (Y) seem to play a major role. Given the weak and transient nature of these contacts, it is more than likely that the much longer IDRs in eukaryotic transcription factors have a similar or even stronger capabilities to alter DNA properties. Biologically, this suggests a new and previously unanticipated function of IDRs as modulators of DNA flexibility and, if several transcription factors need to bind to a promoter, as allosteric elements that provide cooperativity in transcription activation. At a more mechanistic level, our results provide a direct demonstration of dynamic allostery along one-dimensional DNA.

## Methods

### DNA labelling and duplex preparation

DNA constructs (Extended Data Table 1) were synthesized and HPLC-purified by the Keck Biotechnology Resource Laboratory (Yale School of Medicine). For fluorescent labeling with AlexaFluor 488 (donor) and AlexaFluor 594 (acceptor) NHS esters (ThermoFisher), designated thymine or adenine residues were incorporated as C6-dT/dA amino-modified phosphoramidites (Glen Research). For smFRET constructs, labeling positions were chosen to exclude nucleotides previously identified as protected by ComK binding^39^. We followed the labeling protocol by Masoud *et al*.^72^ with the modification of a 1h incubation at 45 °C. Donor and acceptor dyes were conjugated to the forward and reverse ssDNA strands, respectively. Excess dye and unlabeled oligonucleotides were removed by reversed-phase HPLC using a ZORBAX Eclipse Plus C18 column (3.5 µm; Agilent) equilibrated in TEAA buffer (0.1 M, pH 6.5, 5% acetonitrile). A 7–30% acetonitrile gradient over 40 ml was used for separation. Purified fractions were dried (SpeedVac, overnight) and resuspended in 50 µl ddH_2_O. DNA and dye concentrations were determined by UV–Vis spectroscopy using extinction coefficients of AlexaFluor 488 (*ε*_495_= 73,000 M^−1^ cm^−1^) and AlexaFluor 594 (*ε*_590_ = 92,000 M^−1^ cm^−1^). Only samples with a 1:1 dye:ssDNA ratio were used for smFRET. For annealing, 2 pmol donor-labeled ssDNA and 3 pmol acceptor-labeled ssDNA were combined in 50 mM Tris (pH 7.5), 0.1 M NaCl, and 3 mM MgCl_2_, heated to 95 °C, and cooled gradually over 1 h in a PCR cycler. Unlabeled duplexes for cryo-EM were prepared using equimolar forward and reverse strands and the same annealing protocol.

#### Expression and purification of ComK

The sequence of ComK (Uniprot ID: P40396) codon-optimized for expression in *E. coli* (Genscript) (Extended Data Table 3), were inserted into the NcoI and HindIII sites of a pET28b vector, and the protein was expressed in inclusion bodies (IBs) in BL21(DE3) and purified as described previously^19^. Contrary to ComK, the variant ComKΔ expressed in soluble form: *E. coli* cultures were grown in 4 L LB medium supplemented with 50 µg/ml kanamycin and induced at OD_600_ = 0.7 with 0.3 mM IPTG. After 8 h at 37 °C, cells were harvested by centrifugation and resuspended in 30 ml lysis buffer (25 mM Tris-HCl pH 8.0, 0.5 M NaCl, 10 mM imidazole, 100 µM PMSF, 1.2 µg/ml leupeptin, 1 µM pepstatin A). Lysates were sonicated (70% power; 2s on/10 s off; 60 cycles; on ice) and clarified by centrifugation (15,000 rpm, 10 min, 4 °C). The supernatant was filtered (0.45 µm; Sartorius) and applied to a pre-equilibrated 5 ml HisTrap Ni-NTA column (Cytiva) using Buffer A (25 mM Tris-HCl pH 8.0, 0.5 M NaCl, 10 mM imidazole). The column was washed with Buffer A first and subsequently with a high-salt wash buffer (25 mM Tris-HCl pH 8.0, 2 M NaCl, 10 mM imidazole) until the A_280_ signal returned to baseline on an ÄKTA Pure 150 system (Cytiva). The protein was eluted using a 10-min linear gradient at 3 ml/min with Buffer B (25 mM Tris-HCl pH 8.0, 0.5 M NaCl, 500 mM imidazole). The main elution peak was collected, supplemented with 50 mM DTT, incubated on ice for 1 h, and buffer exchanged back into Buffer A using a HiPrep 26/10 desalting column (Cytiva). To enhance purity, the entire Ni-NTA purification procedure was repeated: the sample was reloaded onto a fresh HisTrap column, eluted as before, and again buffer exchanged into Buffer A. His-tag removal was carried out by adding HRV3C protease (0.1 mg/ml) to the eluted protein and incubating for 1 h at room temperature. The cleaved His tag and HRV3C were removed by a final Ni-NTA step, in which untagged protein appeared in the flow-through while tagged species remained bound to the column. The protein was then exchanged into reaction buffer (20 mM Tris-HCl pH 8.0, 150 mM KCl, 5 mM MgCl_2_) using a 5 ml HiTrap desalting column (Cytiva). Protein concentrations were determined using an extinction coefficient of *ε*_280_ = 19,940 M^−1^ cm^−1^, calculated from the ComKΔ amino acid sequence. Due to the presence of four cysteine residues, 10 mM DTT was added to all preparations. Samples were flash-frozen in liquid nitrogen and stored at −80 °C.

#### smFRET experiments

smFRET measurements were carried out in 20 mM Tris pH 7.0, 50 mM L-Arg, 5 mM MgCl_2_, 150 mM KCl, 20 mM DTT and 0.001% Tween-20. Labeled DNA was used at a concentration 30 - 60 pM. The experiments were performed on a MicroTime 200 confocal system (PicoQuant) coupled to an Olympus IX73 microscope body. Donor and acceptor were excited by pulsed interleaved excitation (PIE)^73^ (20 MHz) using a 485 nm diode laser (LDH-D-C-485, PicoQuant) and 594 nm light from a supercontinuum source (Solea, PicoQuant). The excitation beam was directed through a major dichroic (ZT 470-491/594 rpc, Chroma) into a 60×/1.2 NA water objective (Olympus). Samples were measured in custom quartz cuvettes (50 µl). Laser powers were 100 µW (485 nm) and 20 µW (594 nm) at the back aperture. Emission was collected through the same objective, passed through BLP01-488R (Semrock), a 100 µm pinhole, and detected in two channels on SPADs (Excelitas) after spectral separation (T585 LPXR, Chroma) and band-pass filtering (FF03-525/50 and FF02-650/100, Semrock). Photon arrival times were recorded with 16 ps resolution (HydraHarp 400M, PicoQuant). All measurements were performed at 23 °C.

Bursts were identified with a threshold of ≥50 photons for all determinations of fluorescence lifetime decays and FECS functions. For the determination of binding isotherms, a higher threshold of ≥80 photons was used to reduce shot noise. All FRET efficiencies were corrected for instrumental imperfections and the different quantum yields of AlexaFluor488 and AlexaFluor594^74-76^. In addition, molecules without active acceptor were removed using PIE^73^. In addition, for determinations of fluorescence lifetime decays and FECS functions, we only chose molecules that were within the range of ⟨*E*⟩ ± 2*σ*_*E*_ where ⟨*E*⟩ is the mean FRET efficiency of the distribution and *σ*_*E*_ is the width of the FRET histogram after removing molecules with bleached acceptor. This additional constriction allowed us to minimize the population of molecules in which the acceptor dyes bleached during the transit of the molecule throught the confocal volume of the microscope. The FRET efficiency histograms at different concentrations of ComK and its variants were fit with two Gaussian functions to determine the fraction of ComK-bound promoters. Peak positions and widths were fixed to values obtained from ligand-free and saturating conditions, leaving only the Gaussian amplitudes as free parameters. Subpopulation fractions were determined by numerical integration of each Gaussian component. FECS-functions were computed according to Terterov *et al*.^45^. To this end, we first determined all photon pairs separated by a lag time interval (*τ, τ* + Δ*τ*) were Δ*τ* is the time bin (1*μs*) for all bursts. The distribution of the number of photon pairs of the four different types (*N*_*AA*_(*τ*), *N*_*DD*_(*τ*), *N*_*AD*_(*τ*), *N*_*DA*_(*τ*)) were then normalized by the total number of photon pairs *N*_*AA*_(*τ*) + *N*_*DD*_(*τ*) + *N*_*AD*_(*τ*) + *N*_*DA*_(*τ*) = *N*(*τ*), leading to the correlation ratios (*f*_*AA*_ = *N*_*AA*_(*τ*)/*N*(*τ*), *f*_*DD*_ = *N*_*DD*_(*τ*)/*N*(*τ*), *f*_*AD*_ = *N*_*AD*_(*τ*)/*N*(*τ*), *f*_*DA*_ = *N*_*DA*_(*τ*)/*N*(*τ*)). The FRET correlation function (FECS decay) is then given by

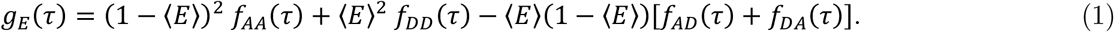

Here, ⟨*E*⟩ is the uncorrected FRET efficiency (proximity ratio) of all bursts used to compute eq. 1. The obtained FRET correlation functions were then fitted with a two-exponential decay with an offset (Fig. 4j, Extended Data Fig. 6a). Examples of the correlation ratios are shown in Extended Data Fig. 6b.

#### Determination of the dimerization state of ComKΔ

The dimerization equilibrium of ComK and ComKΔ was determined using 2fFCS (two-focus fluorescence correlation spectroscopy)^77^. Two overlapping foci were generated in the sample solution by directing two interleaved pulsed laser beams (LDH-D-C-485, PicoQuant, 20MHz each) with orthogonal polarization through a Nomarski prism. The distance between the two resulting foci was determined using the known Stokes radii (*RS*) of the calibration samples: the dye Oregon Green (*RS* = 0.6 nm, 0.36 kDa) and the proteins Aprotinin (*RS* = 1.29 nm, 6.5 kDa), RNaseA (*RS* = 1.73 nm, 13.7 kDa), Carboxy Anhydrase (*RS* = 2.19 nm, 29 kDa), Ovalbumin (*RS* = 2.65 nm, 44 kDa), Conalbumin (*RS* = 3.1 nm, 75 kDa) (Extended Data Fig. 1a,b). For calibration, all proteins were labelled with AlexaFluor488 NHS ester at primary amino groups of lysine residues. The autocorrelation functions of the signal of each focus and the two cross-correlation functions were fit globally with a model that contained one diffusion component and two exponential decay components to account for the triplet dynamics of the dye. The global fits were performed iteratively to minimize the squared difference between the calibration values and the values determined with 2f-FCS. Subsequently, the Stokes radius of ComK and ComKΔ labelled at an introduced cysteine at the terminus was determined in the absence and presence of increasing amounts of unlabeled ComK using the Stokes-Einstein relationship *R*_*S*_ = *k*_*B*_*T*/6*πηD* where *D* is the determined diffusion coefficient, *η* is the viscosity of our buffer (1.333 *cP*), and *k*_*B*_ is Boltzmann’s constant, and *T* is the termperature in Kelvin (276 K) (Extended Data Fig. 1c,d). All measurements were performed at 23°C. Using the calibration samples, the conversion formula from Stokes radii (*R*_*s*_) to molecular weight (*M*) was found to be 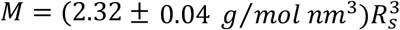. The change of molecular weight as function of the concentration of unlabeled ComK or ComKΔ was fit using the dimerization model *M* = 2*M*_0_*f* + *M*_0_(1 − *f*), where *f* is the fraction of the concentration of labelled ComK or ComKΔ in dimers, as function of the concentration of unlabeled ComK or ComKΔ (*c*)

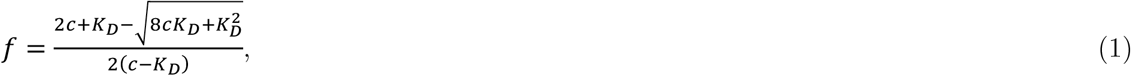

resulting in dissociation constants *K*_*D*_ = 0.53 ± 0.28 *μM* for ComK and *K*_*D*_ = 1.03 ± 0.21 *μM* for ComKΔ. The errors are parameter errors from the fit with equation 1. Here, *M*_0_ = 22.64 *kDa* for ComK and *M*_0_ = 18.16 *kDa* for ComKΔ as computed from the amino acid sequence (Extended Data Table 3).

#### Synchrotron circular dichroism (SRCD) spectroscopy

The SRCD-spectrum of the isolated IDR was determined at the Diamond Light Source (Didcot, UK) at beamline B23. Experiments were performed at a concentration of 67 μM (0.3 mg/ml) in 10mM sodium phosphate pH 8.0, 25mM Na2SO4, 3mM MgSO4. Dynamic light scattering experiments (DLS) carried out at the beamline B23 using the Zetasizer Nano (Malvern) revealed a homogeneous size of ∼3 nm at this concentration, which is larger than the expected size (∼1.3-1.8 nm)^78^, indicating the presence of associations of the peptide. The helical content was computed using CDApps^79^, which resulted in 18% helix, 24% β-strand, 24% turns, 34% disordered.

#### The Pauling model of cooperativity

The Pauling model for the ComK-DNA binding reaction was derived previously^19^ and we provide a brief summary here. Defining *s* = *Kx* for clarity, where *K* is the association constant of a single ComK (or ComKΔ) to a site on the DNA, we obtain the binding polynomial

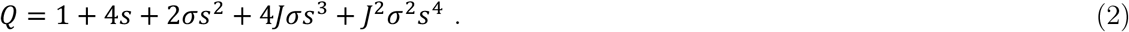

Here, *σ* is the affinity-enhancement for ComK that binds to a box that has already one ComK molecule bound. Similar in spirit, *J* is the affinity-enhancement for ComK that binds to a box if the other box is already saturated with ComK. The fraction of fully bound promoter is therefore given by

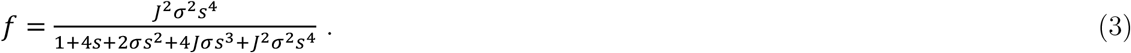

With the same logic, we obtain for the isolated boxes

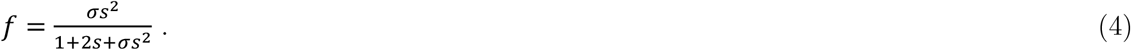

We first fitted the data of the isolated boxes 1 and 2 using eq. 4 (Extended Data Fig. 2) to determine the intra-box cooperativity given by 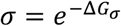. With the value of *σ* determined, we then fitted the data of the promoters with two boxes to determine the inter-box cooperativity 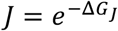. For the determination of these parameters and their error, we globally fitted the binding isotherms from two independent determinations. The error in the parameter is the error of this fit.

#### Sample preparation for cryo-EM

ComKΔ at a concentration of 200 μM in 50 mM Tris pH8, 6 M GdmCl, 10 mM Imidazole, was injected on a HiTrap desalting column (5 ml, GE Healthcare) pre-equilibrated with 0.5 M Arginine-HCl, pH 7.3. The protein containing fractions were then supplemented with 10 mM DTT to prevent the oxidation of the intrinsic cysteine residues of ComKΔ. The sample was concentrated and its concentration was determined by absorbance using *ε*280 = 19440 cm^-1^ M^-1^. The ComKΔ-DNA complexes were formed by quickly mixing 6.43 μl of ComKΔ (560 μM) and 3 μl of DNA (200 μM, 18bp promoter) with 2.5 μl of 10-fold buffer in a total volume of 25ul such that the final buffer composition was 20 mM Tris pH 7.0, 5 mM MgCl2, 150 mM KCl, 3mM DTT, 0.001% Tween-20. Immediately, after mixing, 3.5 μl of this mixture was transferred to UltrAuFoil R 1.2/1.3 300 mesh grids (Quantifoil). The grids were glow discharged before applying the sample, blotted for 2.5 s at 4°C and 100% humidity, and plunge frozen in liquid ethane cooled by liquid nitrogen using a Vitrobot plunger (Thermo Fisher Scientific).

#### Cryo-EM data acquisition

Cryo-EM data sets were collected on a Titan Krios G3i transmission electron microscope (Thermo Fisher Scientific) operated at 300 kV. Movies were recorded in counting mode on a K3 direct detector (Gatan) at the end of BioQuantum energy filter (Gatan) using a slit of 20 eV or 15 eV at a nominal magnification of 105,000×, which corresponds to a physical pixel size of 0.826 Å. The dose rate was set to 17.3 e^-^/pixel/s, and the total exposure time was 2 s, resulting in an accumulated dose of 50.7 e^-^/Å^2^. Each movie was fractionated into 50 frames of 0.04 s. The nominal defocus range was -0.8 to -1.3 μm. Imaging was done using an automated low dose procedure implemented in SerialEM^80^, in which a single image was collected from the center of each hole. Image shift was used to target an array of up to six neighboring holes along the tilt axis (depending on hole array orientation), and stage shift to move between arrays. Beam tilt was adjusted to achieve coma-free alignment when applying image shift.

#### Single particle cryo-EM image processing

Image processing was performed using CryoSPARC software^81^. Movie frames were aligned using patch motion correction, followed patch CTF estimation. 8830 micrographs, which featured a resolution better than 10.0 Å, relative ice thickness lower than 1.4, and full frame motion correction smaller than 100 pixels were selected for further processing. The processing procedure is outlined in Supplementary Fig. 5. An initial particle data set was created by manual picking, followed by 2D classification and automated picking using the newly generated 2D classes as templates. About 1.8M automatically picked particles were extracted, binned 4×4 (98-pixel box size, 3.3 Å/pixel). After 2D classification, we obtained 352,007, 382,195 and 195,939 particles for ComKΔ_4_-DNA, ComKΔ_2_-DNA and free DNA particles, respectively, which were used to determine the ligand distribution shown in Fig. 3a. For ab initio 3D reconstruction of the ComKΔ_4_-DNA particles, we first subjected the data set to additional multiple rounds of 2D classifications. After 3D reconstruction of the ComKΔ_4_-DNA complex with two 3D classes, the higher-resolution 3D class (73,689 particles) was refined using the unbinned data (392 box size) to a final resolution of 6.5 Å. All 3D classes were refined by homogenous refinement using the un-binned data. Final resolutions and angular distributions can be seen in Extended Data Fig. 7. 3D visualization was performed using UCSF Chimera^82^ (Version 1.12) or ChimeraX^83^ (Version 0.92). The 3D-variability analysis was performed according to Punjani and Fleet^43^ using 3 modes, at a filter resolution of 7 Å. The results were then used to generate 20 frames, reporting on the continuous family of 3D structures, derived from particle conformational space (Extended Data Fig. 7).

#### Determination of Hill exponents from cryo-EM distributions

The average number of ligands (*i*) bound to an ensemble of particles is 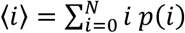 and 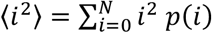 is the second moment of the distribution *p*(*i*) of bound ligands. Here, *N* is the total number of binding sites. Following Wyman^84^, the Hill coefficient is given by

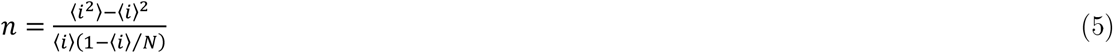

The nominator of eq. 5 is the variance of the observed distribution of bound ligands and the denominator is the variance of the corresponding binomial distribution. We obtain *p*(*i*) from the evaluation of particles in cryo-EM and computed the Hill coefficient with *N* = 4 from the data in Fig. 3a.

#### Determination of distance distributions from smFRET lifetime decays

Fluorescence lifetime decays were constructed from the identified bursts. We computed the decays of the donor 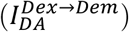 and the acceptor 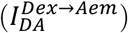 for all molecules with donor and acceptor (*DA*) after exciting the donor (*Dex*). In addition, the decays of the donor in the absence of the acceptor 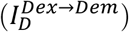 were determined from molecules with inactive acceptor (*D*). Finally, we also determined the decay of the acceptor after directly exciting the acceptor in doubly labeled molecules 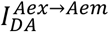. Background correction was performed using the photons not belonging to a burst. We determined the instrumental response functions of the donor excitation in the donor 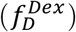 and acceptor channel 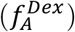 and of the acceptor excitation in the acceptor channel 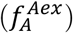 using the back reflection of laser light at the interface between immersion water and quartz coverslip. In the analysis, we first determined the basic emission rate constants of the donor 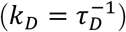 and acceptor 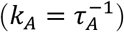 from the decays 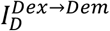 and 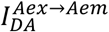 using single-exponential decays convoluted with 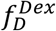 and 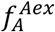, respectively (Extended Data Fig. 4). With the basic emission rate constants determined, we fitted the donor and acceptor decays of the doubly labeled molecules after donor excitation using a convolution with the instrumental response functions 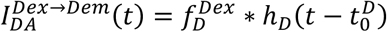 and 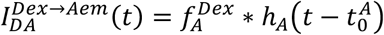 where 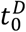 and 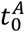 are the time origins of the decays. Here, 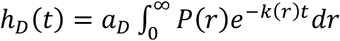 and

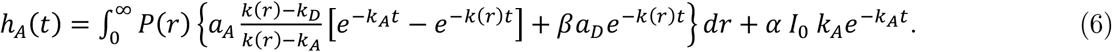

In eq. 6, 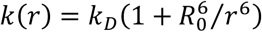 is the total decay rate of the excited state of the donor with the donor-acceptor distance *r* and the Foerster distance *R*_0_ (5.4 nm). Instrumental imperfections due to leakage of donor light into the acceptor channel quantified by *β* (0.049) and the direct excitation of the acceptor dye by the donor laser *α* (0.048) were taken into account. *I*_0_ is the total of all photons emitted from both dyes. The amplitudes *a*_*D*_ and *a*_*A*_ describe the relative brightness of the donor and acceptor decay. The empirical model of the distance distribution is a skewed Gaussian distribution^85^ given by

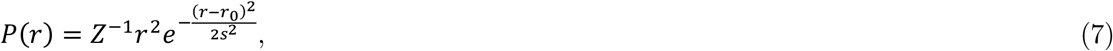

where, *r*_0_ is the position and *s* is the width of the Gaussian, and *Z* is the normalization constant. The global fits of the decays 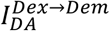 and 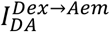 had four free parameters: *r*_0_, *s, a*_*D*_, and *a*_*A*_. The total width *σ* of *P*(*r*) was computed using 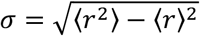. Examples of the lifetime fits with eq. 6-7 are shown in Extended Data Fig. 5.

### Coarse-grained (CG) simulations of the ComK-DNA complex

The CG-Langevin dynamics simulations of the ComK-DNA complex were conducted using OpenMM package v8.2.0..^86^ We used the CG representation and parameters developed for protein-DNA systems by Kapoor *et. al*..^87^ Each amino-acid of the ComK protein was represented with one bead, located at C*α*-position. The DNA was modeled with two beads per base where one bead represents a base and the other represents the sugar-phosphate backbone bead^87^. Bonded interactions included harmonic potentials for all connected beads and angle potentials for DNA backbones. Electrostatic interactions between charged beads were described by screened electrostatic potential with a screening length corresponding to 150 mM ionic strength. The electrostatic potential was truncated at 3.5 nm. Beads representing lysine and arginine were positively charged (+1 elementary charge). Glutamate and aspartate as well as the DNA backbone beads were negatively charged (-1 elementary charge). Short range interactions such as dispersion and hydrophobicity were modeled with potentials and parameters defined in Kapoor *et. al*..^87^ This includes specific nucleotide - amino acid and protein-protein interaction parameters from the HPS-Urry model.^88^ These potentials were truncated at 2.0 nm.

The initial structure of the complex was obtained based on an all-atom model that we fitted into the experimental cryo-EM map of DNA-ComK complex. The model was based on the cryo-EM density map EMD-12260 and the DNA structure with PDB accession code 7NBN. To determine an all-atom structure of wildtype ComK, we predicted the structure using AlphaFold 3.^89^ The predicted all-atom structures of four ComK-molecules, the cryo-EM density map, and the all-atom DNA structure were then imported into UCSF Chimera and the Fit-in-Map functionality was used to perform a rigid-body fit of the ComK models into the cryo-EM density.

To explore possible orientations of the IDRs of ComK, only the IDR regions (residues 158-192) of all four ComK monomers were kept flexible while the remaining beads were fixed to their initial position. In total, we conducted 3 simulations with a length of 10 μs using the Langevin integrator implemented in OpenMM^86^ with a time step of 0.01 ps, a friction coefficient of 1 *ps*^−1^ and a temperature of 300K.

We considered two beads in contact with each other if the distance between them was less then 0.9 nm. This cut-off covers most of the attractive part of the short-range potential and corresponds to the distance between centers of masses of the corresponding groups. We then calculated the probability of quenching by assuming that quenching appears if the center of mass of the quencher, which is the C-terminal residue of each IDR, forms a contact with the sphere representing the dye. Distribution of the positions of the center of mass of the dye bead was estimated from a rotational isomeric state model (RIS) of the dyes attached to the DNA. The determination of the distribution of locations of the dye positions around an attachment site in the DNA was performed earlier in Rosenblum *et al*.^19^. The distribution was fit to a sum of two Gaussian peaks and used to compute the contact probability with the terminal beads of the IDRs.

#### Elastic network model of B-DNA

Ideal double-stranded B-DNA was modelled using a coarse-grained bead-and-spring representation consisting of three beads per nucleotide: phosphate (P), sugar (S), and base (B). The model preserves the double-helical topology while remaining computationally tractable. A total of 1000 bp nucleotides base pairs were used to eliminate finite-length effects. Backbone connectivity was enforced by harmonic springs between consecutive phosphate and sugar beads along each strand. Base–base and base–backbone interactions were introduced using a distance-based cut-off to represent non-bonded contacts. All interactions were modelled as harmonic springs. The system was described by an elastic network model (ENM) in which interacting bead pairs (*i, j*) were connected by harmonic springs with the arbitrarily chosen spring constant *γ*_*i,j*_ = 1 *k*_*B*_*T*/Å^+^ with *k*_*B*_*T* = 1. With **r**_*i*_ = (*x*_*i*_, *y*_*i*_, *z*_*i*_) being the position vector of bead *i*, the total potential is given by

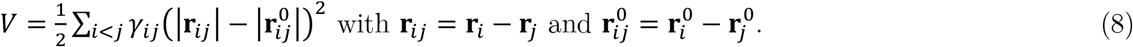

Here, 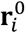 is the equilibrium position of bead *i*. The elastic energy of a deformed DNA can also be written as 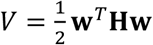 with the 3*N* × 3*N* Hessian matrix **H**, and **w** is the column vector of bead displacements. Here, *N* is the total number of beads. We determined the eigenvalues (*λ*^(*m*)^) and eigenvectors (**u**^(*m*)^) of **H** for each mode *m* by solving the eigenvalue problem **H u**^(*m*)^ = *λ*^(*m*)^**u**^(*m*)^ with *m* = {1,2,3, …,3*N*}. We then determine the equilibrium unit vector between bead *q* in base pairs *k* and *l* where *q* = {1, 2, …,6} is the local summation index for the beads in a base pair, which is given by 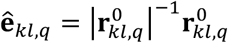 . We then compute the fluctuations in distance *δd*_*kl*_ between base pair *k* and *l* via^51^

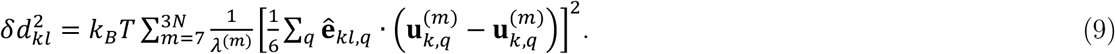

The distance fluctuations were computed for a window of 9 bp, i.e., *l* = *k* + 8, where the window was moved over the total length of the DNA in steps of 2 bp. This results in a profile of distance fluctuations along the DNA. To minimize local effects from the intrinsic helicity of the DNA, we also averaged these profiles over different positions from *k* = 200 to *k* = 792 (*l* = 800) in steps of 5bp, which provides the averaged profile 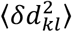 . Finally, to study the effect of a “perturbation”, we increased the spring constants of the bonds within a region (box) of nine consecutive base pairs 3-fold and 5-fold and computed 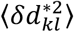, which is the profile of distance fluctuations of a perturbed DNA. In Fig. 5g, we plot the difference in fluctuations 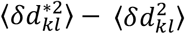.

## Data availability

Cryo-EM maps for the ComKΔ-DNA complex have been deposited in the Electron Microscopy Data Bank (EMD-56533).

## Code availability

The single-molecule analysis code is accessible as a custom WSTP add-on for Mathematica (Wolfram Research) at https://schuler.bioc.uzh.ch/programs/.

## Acknowledgments

We thank Haim Rozenberg for critical discussions on the cryo-EM data, Donna Matzov for her advice with Cryo-EM data processing, and Harry Greenblatt for server configuration and installation support, and Diamond Light Source Ltd for beamtime on B23 beamline (SM35669 and SM37175). This work was supported by a grant of the European Research Council (Grant No. 864578) to HH.

