## Supplementary Information for "Protein disorder controls allostery in DNA"

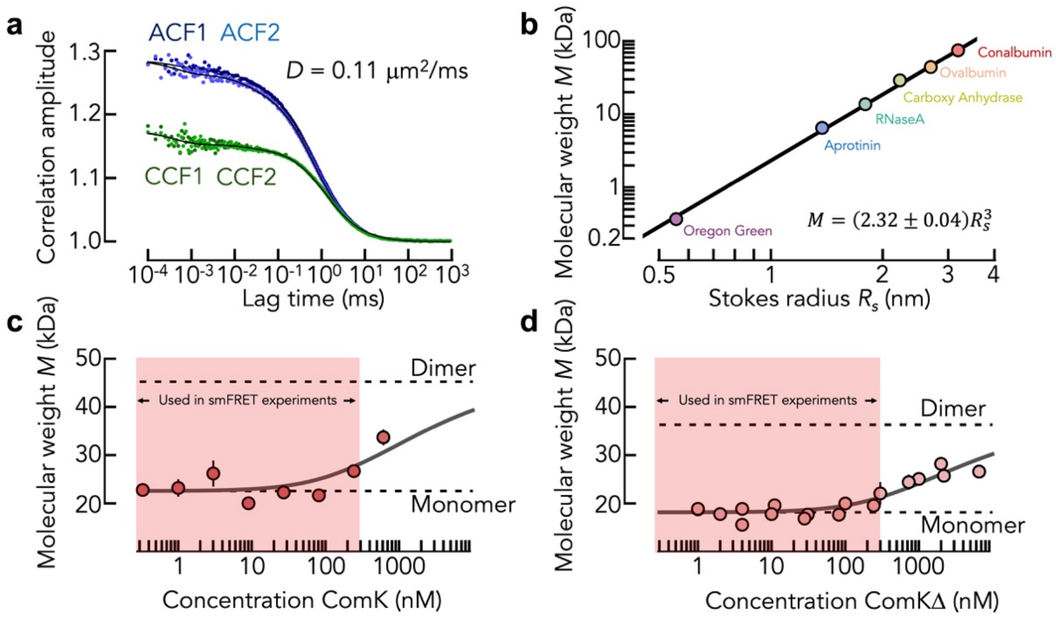

**Extended Data Fig. 1 Determination of the oligomerization state of ComK and ComKΔ using 2fFCS**. **(a)** Examples of auto- (blue) and cross- (green) intensity correlation functions of ComKΔ, labeled with AlexaFluor488. Black solid lines are fits with a model of a decay accounting for diffusion through the two confocal volumes, and two exponential decays to account for the triplet dynamics of the dye. The resulting diffusion coefficient is indicated. **(b)** Scaling between molecular weight and the Stokes radii of the calibration samples. The functional form of the scaling is indicated. **(c, d)** Molecular weight of labeled ComK (c) and ComKΔ (d) in the presence of increasing amounts of unlabeled ComK. The solid line is a fit with eq. 1 (Methods) and dashed lines indicate the molecular weight of monomer and dimers. Error bars indicate the standard deviation of three technical replicates. **(d)** Molecular weight of labeled ComKΔ in the presence of increasing amounts of unlabeled ComK. The solid line is a fit with eq. 1 (Methods). Error bars is the standard deviation of three technical replicates.

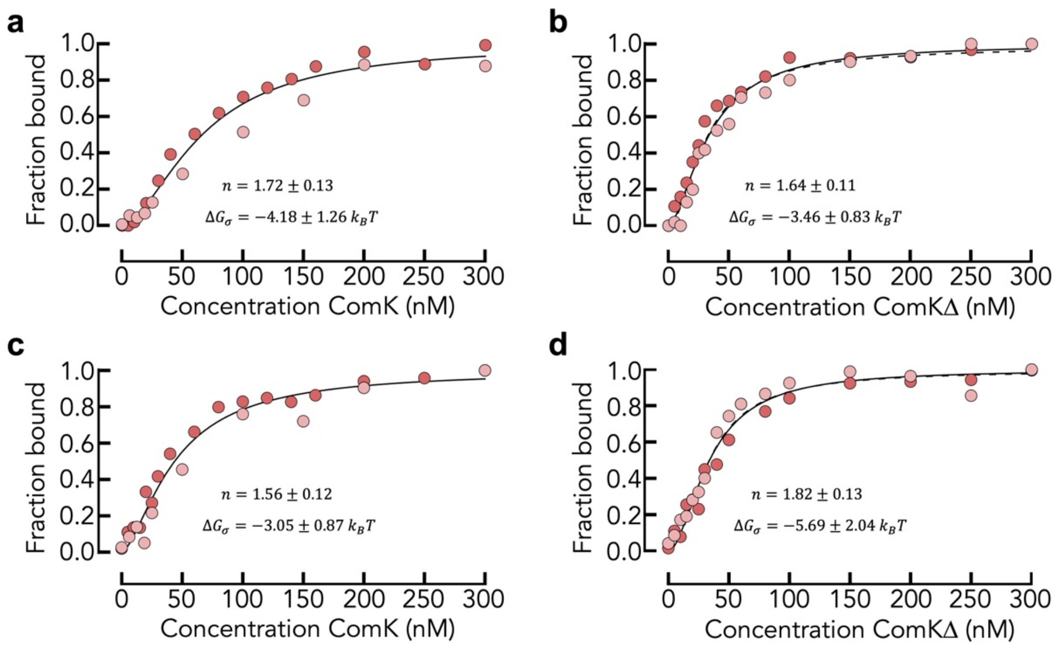

**Extended Data Fig. 2 Binding isotherms to single-box promoters determined with smFRET. (a, b)** Fraction of bound promoter (single-box 1) with increasing concentration of ComK (a) and ComKΔ (b). **(c, d)** Fraction of bound promoter (single-box 1) with increasing concentration of ComK (c) and ComKΔ (d). Dark and light red are two independent experiments. The solid lines are global fits with the Hill equation, and the dashed lines are global fits with a two-site Pauling model. The Hill exponent and the free energy of coupling between the two sites is indicated. The error in the parameters is the error of the global fit.

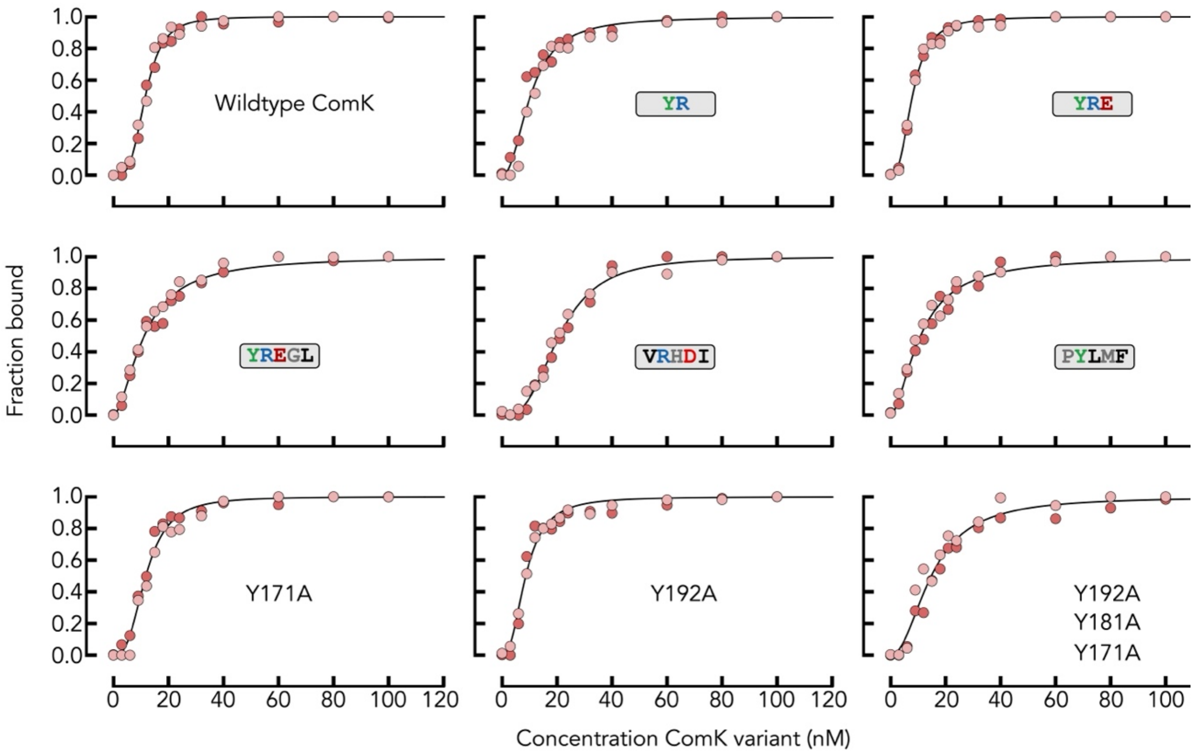

**Extended Data Fig. 3 Binding isotherms of ComK with modified IDRs to the two-box promoter with an 18bp spacer smFRET.** The fraction of bound promoter with increasing concentration of the modified ComK is shown. Dark and light red are two independent experiments. The solid lines are global fits with the Hill equation. The modification of the IDR is indicated.

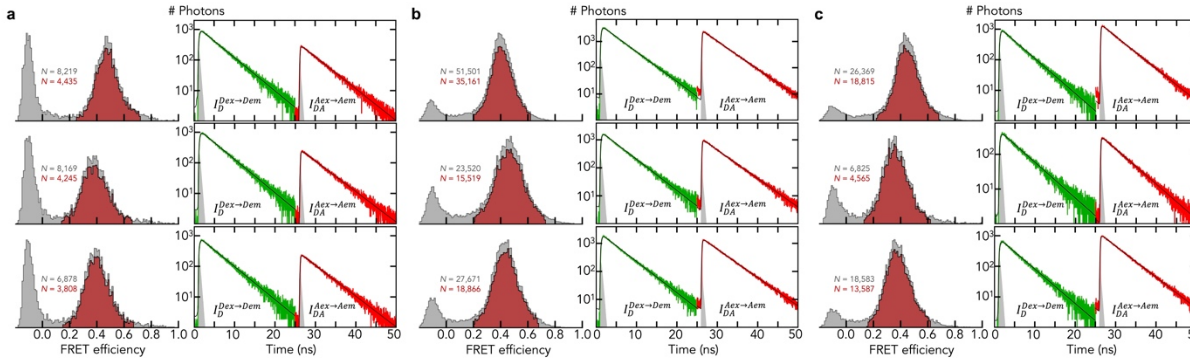

**Extended Data Fig. 4 Determination of the intrinsic fluorescence lifetimes of the donor and acceptor dye in the absence of FRET. (a, left)** FRET histograms of the 18bp-promoter labeled in Box1 in the absence (top) and presence of ComK (middle) and ComKΔ (bottom). The gray area indicates all molecules whereas the red area indicates molecules after removing donor-only and bleached molecules. The molecule numbers are indicated. **(a, right)** Fluorescence lifetime decays of donor (green) in the absence of the acceptor and acceptor (red) after direct excitation of the acceptor together with single-exponential fits of the decays. (b, c) Same as a but for an 18bp-promoter with labeled spacer (b) and an 18bp-promoter with a labeled box2 (c). The concentration of ComK and ComKΔ was 100 nM.

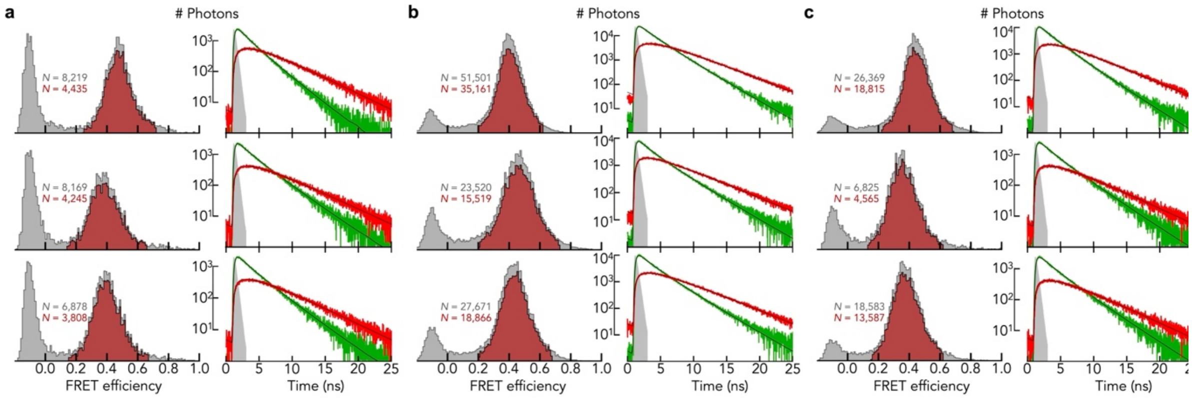

**Extended Data Fig. 5 Determination of the distance distribution in the 18bp-promoter using fluorescence lifetime experiments. (a, left)** FRET histograms of the 18bp-promoter labeled in Box1 in the absence (top) and presence of ComK (middle) and ComKΔ (bottom). The gray area indicates all molecules whereas the red area indicates molecules after removing donor-only and bleached molecules. The molecule numbers are indicated. **(a, right)** Fluorescence lifetime decays of donor (green) and acceptor (red) for doubly-labeled molecules after donor excitation. Solid lines are global fits with eq. 5-6. **(b, c)** Same as a but for an 18bp-promoter with labeled spacer (b) and a labeled box2 (c). The concentration of ComK and ComKΔ was 100 nM.

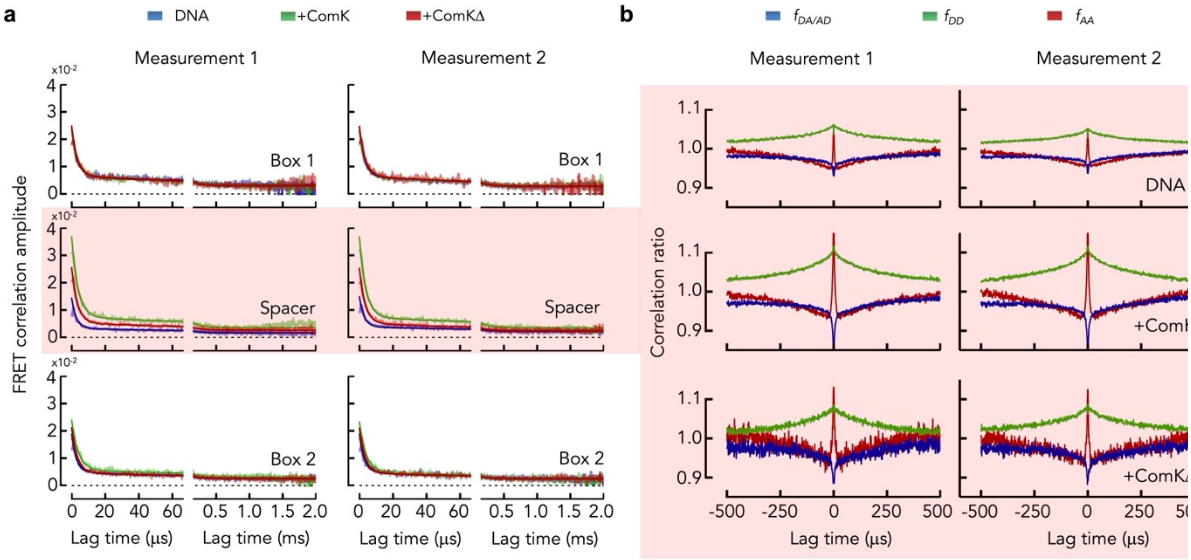

**Extended Data Fig. 6 Examples of FECS decays and correlation ratios for the two-box promoter with an 18bp spacer. (a)** FECS decays of two independent measurements (left and right) in the absence and presence of 100 nM ComK and ComKΔ (color indicated). **(b)** Correlation ratios ($f_{AA} = N_{AA}(\tau)/N(\tau)$, $f_{DD} = N_{DD}(\tau)/N(\tau)$, $f_{AD} = N_{AD}(\tau)/N(\tau)$, $f_{DA} = N_{DA}(\tau)/N(\tau)$) as function of the lag time.

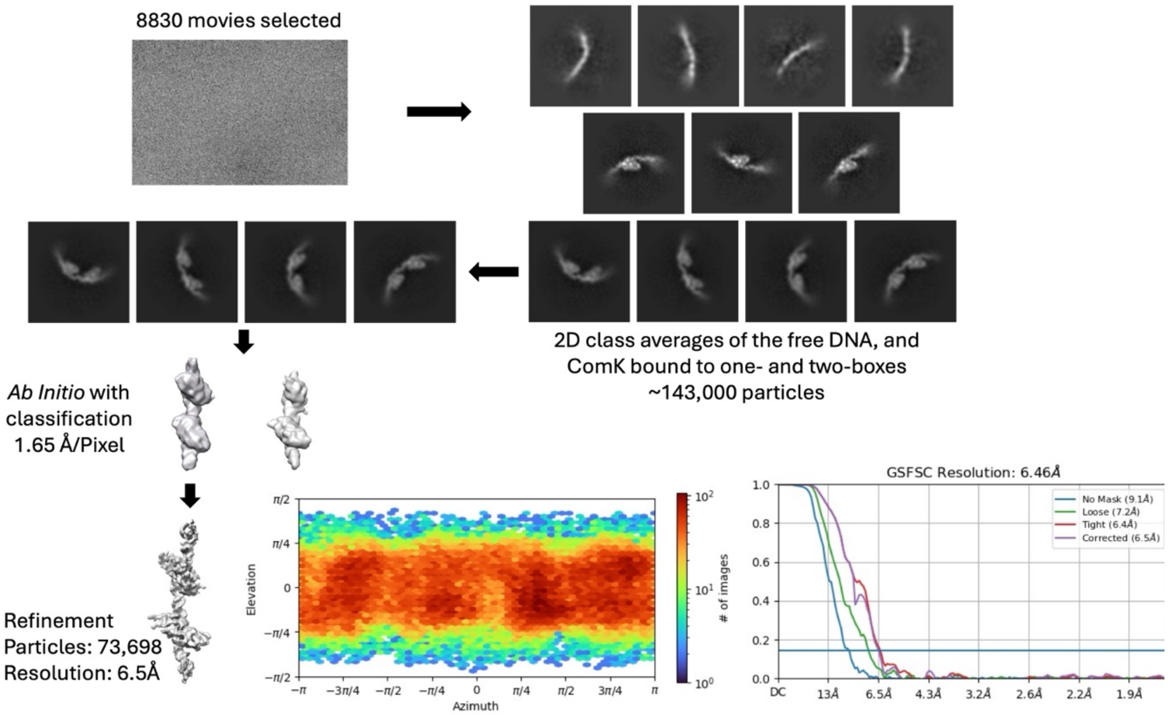

**Extended Data Fig. 7. Scheme for single particle reconstruction of the 18 bp data set with ComKΔ.** The details of the process are described in the Methods section. Briefly, a mixed particle data set was first extracted from the micrographs, containing both free and ComK-bound DNA complexes. This mixed data set was cleaned and separated into two data sets, ComK-DNA and free DNA, by iterative 2D classification. Processing, followed by ab initio 3D reconstruction and classification, and refinement of the best 3D class. Angular distribution and FSC curves are presented for the final 3D map.

**Extended Data Table 1. List of labelled DNA for smFRET ComK-binding experiments.** The two binding boxes are highlighted in cyan. Donor and acceptor attachment sites are shown in green and red, respectively. For the constructs of the isolated boxes, we added a GC pair at the termini to prevent helix fraying. The construct denoted by *Double-box 18bp* is identical to the natural *comG* promoter.

| Promoter | *bp* | Sequence |
|---|---|---|
| ***Single-box 1*** | 46 | CTTTTTTCTTGCCAGAAAGAATTGGTTTT**T**CAGCATATAACATCTC<br>GAAAAAAGAACGGTCT**T**TCTTAACCAAAAAGTCGTATATTGTAGAG |
| ***Single-box 2*** | 43 | CATATAACATCTCACAAAATCACGTTT**T**CCCTGTTTGATTACC<br>GTATATTGTAGAG**T**GTTTTAGTGCAAAAGGGACAAACTAATGG |
| ***Double-box 8bp*** | 78 | GAAAGTCTTTTTTCTTGCCAGAAAGAATTGGTTTT**T**TATATGTGAAAATCACGTTTTCCCTGTTTGATTACCTTTTCT<br>CTTTCAGAAAAAAGAACGGTCT**T**TCTTAACCAAAAAATATACACTTTTAGTGCAAAAGGGACAAACTAATGGAAAAGA |
| ***Double-box 14bp*** | 84 | GAAAGTCTTTTTTCTTGCCAGAAAGAATTGGTTTT**T**CAGCATATAACATCAAAATCACGTTTTCCCTGTTTGATTACCTTTTCT<br>CTTTCAGAAAAAAGAACGGTCT**T**TCTTAACCAAAAAGTCGTATATTGTAGTTTTAGTGCAAAAGGGACAAACTAATGGAAAAGA |
| ***Double-box 18bp*** | 93 | GAAAGTCTTTTTTCTTGCCAGAAAGAATTGGTTTT**T**CAGCATATAACATCTCACAAAATCACGTTTTCCCTGTTTGATTACCTTTTCT<br>CTTTCAGAAAAAAGAACGGTCT**T**TCTTAACCAAAAAGTCGTATATTGTAGAGTGTTTTAGTGCAAAAGGGACAAACTAATGGAAAAGA |
| ***Double-box 24bp*** | 94 | GAAAGTCTTTTTTCTTGCCAGAAAGAATTGGTTTT**T**CAGCATATAACATCTCACCAGCATAAAATCACGTTTTCCCTGTTTGATTACCTTTTCT<br>CTTTCAGAAAAAAGAACGGTCT**T**TCTTAACCAAAAAGTCGTATATTGTAGAGTGGTCGTATTTTAGTGCAAAAGGGACAAACTAATGGAAAAGA |
| ***Double-box 31bp*** | 101 | GAAAGTCTTTTTTCTTGCCAGAAAGAATTGGTTTT**T**TACTAGTCATTTAGTACCATTAAATATCATTAAAATCACGTTTTCCCTGTTTGATTACCT<br>TTTCT<br>CTTTCAGAAAAAAGAACGGTCT**T**TCTTAACCAAAAAATGATCAGTAAATCATGGTAATTTATAGTAATTTTAGTGCAAAAGGGACAAACTAATGGA<br>AAAGA |
| ***Designed box*** | 89 | CTTGCCAGAATCAGCA**T**ATAACATCTCACTCAGCATATAACATCTCAC**T**CAGCATATAACATCTCACAAAATCACGTTT**T**CCCTGTTTG<br>GAACGGTCTTAGTCGTATATTGTAGAGTGAG**T**CGTATATTGTAGAGTGAGTCGTATATTGTAGAG**T**GTTTTAGTGCAAAAGGGACAAAC |
| ***Designed sp3*** | 89 | CTTGCCAGAATCAGCA**T**ATAACATCTCACTCAGCATATAACATCTCAC**T**CAGCATATAACATCTCACAAAATCACGTTT**T**CCCTGTTTG<br>GAACGGTCTTAGTCGTATATTGTAGAGTGAG**T**CGTATATTGTAGAGTGAGTCGTATATTGTAGAG**T**GTTTTAGTGCAAAAGGGACAAAC |
| ***Designed sp2*** | 89 | CTTGCCAGAATCAGCA**T**ATAACATCTCACTCAGCATATAACATCTCAC**T**CAGCATATAACATCTCACAAAATCACGTTT**T**CCCTGTTTG<br>GAACGGTCTTAGTCGTATATTGTAGAGTGAG**T**CGTATATTGTAGAGTGAGTCGTATATTGTAGAG**T**GTTTTAGTGCAAAAGGGACAAAC |
| ***Designed sp1*** | 89 | CTTGCCAGAATCAGCA**T**ATAACATCTCACTCAGCATATAACATCTCAC**T**CAGCATATAACATCTCACAAAATCACGTTT**T**CCCTGTTTG<br>GAACGGTCTTAGTCGTATATTGTAGAGTGAG**T**CGTATATTGTAGAGTGAGTCGTATATTGTAGAG**T**GTTTTAGTGCAAAAGGGACAAAC |

**Extended Data Table 2. Fitting parameters obtained with the mechanistic binding model and with the Hill equation.** Errors result from global fit of at least two independent experiments. To increase the robustness of the fit parameters in our mechanistic binding model. Fixed parameters are indicated by (*) .

| Promoter | Variant | $\Delta g_K$ ($k_BT$) | $-\Delta g_\sigma$ ($k_BT$) | $-\Delta g_J$ ($k_BT$) | $K_{Hill}$ (nM) | $n$ |
|---|---|---|---|---|---|---|
| *Single box 1* | *ComK* | 6.14 ± 0.73 | 4.18 ± 1.26 | - | 66.2 ± 3.2 | 1.72 ± 0.13 |
| *Single box 2* | *ComK* | 5.05 ± 0.54 | 3.05 ± 0.87 | - | 43.7 ± 2.4 | 1.56 ± 0.12 |
| *Single box Global* | *ComK* | 5.23 ± 0.47 | 3.02 ± 0.75 | - | 53.4 ± 2.5 | 1.58 ± 0.10 |
| *Design 8bp* | *ComK* | 6.21 ± 0.53 | 3.02* | 4.23 ± 0.98 | 14.3 ± 0.4 | 3.60 ± 0.34 |
| *Design 14bp* | *ComK* | 5.55 ± 0.27 | 3.02* | 1.86 ± 0.40 | 30.1 ± 0.9 | 2.41 ± 0.18 |
| *Design 18bp* | *ComK* | 6.73 ± 0.52 | 3.02* | 5.61 ± 0.99 | 11.6 ±0.2 | 3.68 ± 0.20 |
| *Design 24bp* | *ComK* | 5.40 ± 0.16 | 3.02* | 1.73 ± 0.23 | 28.1 ± 0.4 | 2.44 ± 0.09 |
| *Design 31bp* | *ComK* | 6.16 ± 0.24 | 3.02* | 3.12 ± 0.41 | 25.2 ± 0.6 | 2.82 ± 0.18 |
| *Single box 1* | *ComKΔ* | 5.05 ± 0.50 | 3.46 ± 0.83 | - | 33.6± 1.3 | 1.64 ± 0.11 |
| *Single box 2* | *ComKΔ* | 6.31 ± 1.09 | 5.69 ± 2.04 | - | 33.9 ± 1.3 | 1.82 ± 0.13 |
| *Single box Global* | *ComKΔ* | 5.53 ± 0.45 | 4.28 ± 0.79 | - | 33.7 ± 0.9 | 1.72 ± 0.08 |
| *Design 8bp* | *ComKΔ* | 4.44 ± 0.40 | 4.28* | 0.00 ± 0.67 | 19.0 ± 0.5 | 2.06 ± 0.12 |
| *Design 14bp* | *ComKΔ* | 3.88 ± 0.41 | 4.28* | 0.00 ± 1.61 | 10.6 ± 0.3 | 1.64 ± 0.08 |
| *Design 18bp* | *ComKΔ* | 4.23 ± 0.18 | 4.28* | 0.26 ± 0.21 | 11.3 ±0.3 | 1.82 ± 0.09 |
| *Design 24bp* | *ComKΔ* | 4.03 ± 0.39 | 4.28* | 0.00 ± 0.80 | 12.7 ± 0.3 | 1.71 ± 0.08 |
| *Design 31bp* | *ComKΔ* | 4.42 ± 0.35 | 4.28* | 0.00 ± 0.41 | 18.7 ± 0.6 | 1.98 ± 0.13 |
| *Design 18bp* | *YR* | - | - | - | 10.3 ± 0.5 | 2.16 ± 0.19 |
| *Design 18bp* | *YRE* | - | - | - | 7.6 ± 0.2 | 2.68 ± 0.11 |
| *Design 18bp* | *YREGL* | - | - | - | 11.2 ± 0.3 | 1.73 ± 0.09 |
| *Design 18bp* | *VRHDI* | - | - | - | 20.8 ±0.4 | 2.88 ± 0.15 |
| *Design 18bp* | *PYLMF* | - | - | - | 10.9 ± 0.4 | 1.70 ± 0.10 |
| *Design 18bp* | *Y171A* | - | - | - | 11.8 ± 0.3 | 2.87 ± 0.21 |
| *Design 18bp* | *Y192A* | - | - | - | 8.6 ± 0.3 | 2.45 ± 0.18 |
| *Design 18bp* | *Y171A/Y181A/Y192A* | - | - | - | 14.8 ± 0.6 | 2.10± 0.18 |

**Extended Data Table 3. Amino acid sequences of the ComK constructs used in this study.** The codon optimized construct contained a His6-tag (red) and an HRV3C cleavage site (cyan) used for all smFRET and cryo-EM experiments (top). The natural cysteine residues of ComK (yellow) are not surface exposed. For determination of the oligomerization state using 2fFCS, we also created constructs with a surface-exposed additional cysteine at the N-terminus (bottom). Vertical line indicates the HRV3C cleavage position.

<table>
<tr><th>Variant</th><th>Amino acid sequence</th></tr>
<tr><td>ComK</td><td>MGSSHHHHHHSGSGSAGLEVLFQ|GPGMSQKTDAPLESYEVNGATIAVLPEEIDGKICSKIIEKDCVFYVNMKPLQIVDRSCRFFGSSYAGRKAGTYEVTKI<br>SHKPPIMVDPSNQIFLFPTLSSTRPQCGWISHVHVKEFKATEFDDTEVTFSNGKTMELPISYNSFENQVYRTAWLRTKFQDRIDHRVPKRQEFMLYPKEERT<br>KMIYDFILRELGERY</td></tr>
<tr><td>ComKΔ</td><td>MGSSHHHHHHSGSGSAGLEVLFQ|GPGMSQKTDAPLESYEVNGATIAVLPEEIDGKICSKIIEKDCVFYVNMKPLQIVDRSCRFFGSSYAGRKAGTYEVTKI<br>SHKPPIMVDPSNQIFLFPTLSSTRPQCGWISHVHVKEFKATEFDDTEVTFSNGKTMELPISYNSFENQVYRTAWLRTKFQDR</td></tr>
<tr><td>YR</td><td>MGSSHHHHHHSGSGSAGLEVLFQ|GPGMSQKTDAPLESYEVNGATIAVLPEEIDGKICSKIIEKDCVFYVNMKPLQIVDRSCRFFGSSYAGRKAGTYEVTKI<br>SHKPPIMVDPSNQIFLFPTLSSTRPQCGWISHVHVKEFKATEFDDTEVTFSNGKTMELPISYNSFENQVYRTAWLRTKFQDRIDHRVPKRQEFMLYPKEERT<br>KMIYDFILRELGE</td></tr>
<tr><td>YRE</td><td>MGSSHHHHHHSGSGSAGLEVLFQ|GPGMSQKTDAPLESYEVNGATIAVLPEEIDGKICSKIIEKDCVFYVNMKPLQIVDRSCRFFGSSYAGRKAGTYEVTKI<br>SHKPPIMVDPSNQIFLFPTLSSTRPQCGWISHVHVKEFKATEFDDTEVTFSNGKTMELPISYNSFENQVYRTAWLRTKFQDRIDHRVPKRQEFMLYPKEERT<br>KMIYDFILRELG</td></tr>
<tr><td>YREGL</td><td>MGSSHHHHHHSGSGSAGLEVLFQ|GPGMSQKTDAPLESYEVNGATIAVLPEEIDGKICSKIIEKDCVFYVNMKPLQIVDRSCRFFGSSYAGRKAGTYEVTKI<br>SHKPPIMVDPSNQIFLFPTLSSTRPQCGWISHVHVKEFKATEFDDTEVTFSNGKTMELPISYNSFENQVYRTAWLRTKFQDRIDHRVPKRQEFMLYPKEERT<br>KMIYDFILRE</td></tr>
<tr><td>PYLMF</td><td>MGSSHHHHHHSGSGSAGLEVLFQ|GPGMSQKTDAPLESYEVNGATIAVLPEEIDGKICSKIIEKDCVFYVNMKPLQIVDRSCRFFGSSYAGRKAGTYEVTKI<br>SHKPPIMVDPSNQIFLFPTLSSTRPQCGWISHVHVKEFKATEFDDTEVTFSNGKTMELPISYNSFENQVYRTAWLRTKFQDRIDHRVPKRQEKEERTKMIYD<br>FILRELGERY</td></tr>
<tr><td>VRHDI</td><td>MGSSHHHHHHSGSGSAGLEVLFQ|GPGMSQKTDAPLESYEVNGATIAVLPEEIDGKICSKIIEKDCVFYVNMKPLQIVDRSCRFFGSSYAGRKAGTYEVTKI<br>SHKPPIMVDPSNQIFLFPTLSSTRPQCGWISHVHVKEFKATEFDDTEVTFSNGKTMELPISYNSFENQVYRTAWLRTKFQDRPKRQEFMLYPKEERTKMIYD<br>FILRELGERY</td></tr>
<tr><td>Y171A</td><td>MGSSHHHHHHSGSGSAGLEVLFQ|GPGMSQKTDAPLESYEVNGATIAVLPEEIDGKICSKIIEKDCVFYVNMKPLQIVDRSCRFFGSSYAGRKAGTYEVTKI<br>SHKPPIMVDPSNQIFLFPTLSSTRPQCGWISHVHVKEFKATEFDDTEVTFSNGKTMELPISYNSFENQVYRTAWLRTKFQDRIDHRVPKRQEFML**A**PKEERT<br>KMIYDFILRELGERY</td></tr>
<tr><td>Y192A</td><td>MGSSHHHHHHSGSGSAGLEVLFQ|GPGMSQKTDAPLESYEVNGATIAVLPEEIDGKICSKIIEKDCVFYVNMKPLQIVDRSCRFFGSSYAGRKAGTYEVTKI<br>SHKPPIMVDPSNQIFLFPTLSSTRPQCGWISHVHVKEFKATEFDDTEVTFSNGKTMELPISYNSFENQVYRTAWLRTKFQDRIDHRVPKRQEFMLYPKEERT<br>KMIYDFILRELGER**A**</td></tr>
<tr><td>Y171A/Y181A<br>/Y192A</td><td>MGSSHHHHHHSGSGSAGLEVLFQ|GPGMSQKTDAPLESYEVNGATIAVLPEEIDGKICSKIIEKDCVFYVNMKPLQIVDRSCRFFGSSYAGRKAGTYEVTKI<br>SHKPPIMVDPSNQIFLFPTLSSTRPQCGWISHVHVKEFKATEFDDTEVTFSNGKTMELPISYNSFENQVYRTAWLRTKFQDRIDHRVPKRQEFML**A**PKEERT<br>KMI**A**DFILRELGER**A**</td></tr>
<tr><td>ComK for<br>2fFCS</td><td>MGSSHHHHHHSGSGSAGLEVLFQ|GP**C**MSQKTDAPLESYEVNGATIAVLPEEIDGKI**C**SKIIEKD**C**VFYVNMKPLQIVDRS**C**RFFGSSYAGRKAGTYEVTKI<br>SHKPPIMVDPSNQIFLFPTLSSTRPQ**C**GWISHVHVKEFKATEFDDTEVTFSNGKTMELPISYNSFENQVYRTAWLRTKFQDRIDHRVPKRQEFMLYPKEERT<br>KMIYDFILRELGERY</td></tr>
<tr><td>ComKΔ for<br>2fFCS</td><td>MGSSHHHHHHSGSGSAGLEVLFQ|GP**C**MSQKTDAPLESYEVNGATIAVLPEEIDGKI**C**SKIIEKD**C**VFYVNMKPLQIVDRS**C**RFFGSSYAGRKAGTYEVTKI<br>SHKPPIMVDPSNQIFLFPTLSSTRPQ**C**GWISHVHVKEFKATEFDDTEVTFSNGKTMELPISYNSFENQVYRTAWLRTKFQDR</td></tr>
</table>

**Extended Data Table 4. Parameters of fits of the fluorescence lifetime decays.** Parameters are shown for independent experiment.

| Construct | $\tau_D$ (ns) | $\tau_A$ (ns) | $I_0$ | $a_D$ | $a_A$ | $s$ (nm) | $r_0$ (nm) |
|---|---|---|---|---|---|---|---|
| Construct: Box1-Sp-Box2 Fig. 4i | | | | | | | |
| Box1 | 4.00 | 4.17 | 10,825 | 3,834 | 1,666 | 0.593 | 5.358 |
| | 3.89 | 4.17 | 35,194 | 13,218 | 5,159 | 0.606 | 5.283 |
| Box1+ComK | 3.98 | 4.34 | 10,298 | 3,405 | 1,395 | 0.812 | 5.609 |
| | 3.96 | 4.35 | 18,633 | 5,917 | 2,617 | 0.776 | 5.672 |
| Box1+ComKΔ | 4.02 | 4.30 | 9,061 | 3,239 | 1,229 | 0.753 | 5.540 |
| | 4.06 | 4.36 | 6,019 | 2,104 | 808 | 0.795 | 5.596 |
| Sp | 3.82 | 4.06 | 108,828 | 32,646 | 17,082 | 0.607 | 5.702 |
| | 3.86 | 4.12 | 67,311 | 20,307 | 11,033 | 0.641 | 5.731 |
| Sp+ComK | 3.90 | 4.25 | 37,442 | 11,520 | 5,915 | 1.045 | 5.297 |
| | 3.88 | 4.29 | 41,300 | 12,616 | 6,540 | 1.070 | 5.320 |
| Sp+ComKΔ | 3.89 | 4.27 | 46,647 | 13,767 | 7,457 | 0.853 | 5.586 |
| | 3.92 | 4.26 | 48,096 | 14,345 | 7,482 | 0.858 | 5.560 |
| Box2 | 3.80 | 4.11 | 48,163 | 15,374 | 7,820 | 0.635 | 5.556 |
| | 3.85 | 4.13 | 22,410 | 7,043 | 3,622 | 0.624 | 5.515 |
| Box2+ComK | 4.03 | 4.32 | 12,572 | 4,137 | 1,576 | 0.903 | 5.654 |
| | 4.22 | 4.30 | 11,152 | 3,565 | 1,381 | 0.728 | 5.623 |
| Box2+ComKΔ | 3.85 | 4.27 | 33,828 | 10,133 | 5,110 | 0.805 | 5.815 |
| | 3.90 | 4.27 | 21,982 | 6,645 | 3,203 | 0.818 | 5.777 |
| Construct: Sp1-Sp2-Sp3-Box Fig. 5a | | | | | | | |
| Box | 4.00 | 4.08 | 43,073 | 9,617 | 11,532 | 0.995 | 5.349 |
| | 4.06 | 4.03 | 12,081 | 2,810 | 3.128 | 0.844 | 5.326 |
| | 3.88 | 4.27 | 38,247 | 9,410 | 9,241 | 0.825 | 5.312 |
| | 3.91 | 4.30 | 41,382 | 9,893 | 10,023 | 0.876 | 5.301 |
| Box+ComK | 4.07 | 4.29 | 19,521 | 4,488 | 4,911 | 1.245 | 5.410 |
| | 4.07 | 4.38 | 15,370 | 3,651 | 3,596 | 1.190 | 5.414 |
| Box+ComKΔ | 3.89 | 4.42 | 32,394 | 7,848 | 7,663 | 1.091 | 5.441 |
| | 3.92 | 4.44 | 37,450 | 9,253 | 8,598 | 1.093 | 5.390 |
| Sp1 | 4.15 | 4.14 | 6,763 | 1,819 | 1,659 | 0.854 | 4.644 |
| | 4.13 | 4.07 | 12,986 | 3,500 | 3,112 | 1.054 | 4.299 |
| | 3.77 | 4.42 | 17,843 | 4,607 | 4,213 | 0.867 | 4.642 |
| | 3.87 | 4.50 | 18,195 | 4,584 | 4,102 | 0.944 | 4.507 |
| Sp1+ComK | 4.28 | 4.24 | 3,731 | 1,266 | 826 | 0.946 | 4.387 |
| | 4.29 | 4.18 | 8,673 | 2,368 | 1,983 | 1.004 | 4.341 |
| Sp1+ComKΔ | 3.86 | 4.33 | 22,648 | 6,494 | 5,229 | 0.877 | 4.580 |
| | 3.89 | 4.38 | 13,961 | 3,353 | 3,255 | 0.938 | 4.601 |
| Sp2 | 4.11 | 4.16 | 10,052 | 2,338 | 2,638 | 0.910 | 5.568 |
| | 4.15 | 4.07 | 8,502 | 2,090 | 2,206 | 0.870 | 5.463 |
| | 3.89 | 4.34 | 35,148 | 8,066 | 8,549 | 0.933 | 5.511 |
| | 3.86 | 4.32 | 41,559 | 9,998 | 10,284 | 0.956 | 5.553 |
| Sp2+ComK | 4.29 | 4.20 | 8,300 | 2,009 | 20,37 | 0.928 | 5.368 |
| | 4.24 | 4.18 | 6,386 | 1,820 | 1,497 | 0.904 | 5.229 |
| Sp2+ComKΔ | 3.89 | 4.32 | 41,354 | 9,854 | 9,918 | 0.899 | 5.505 |
| | 3.89 | 4.31 | 42,191 | 9,580 | 10,409 | 0.912 | 5.555 |
| Sp3 | 4.01 | 4.11 | 18,407 | 4,048 | 5,169 | 1.010 | 5.945 |
| | 4.05 | 4.08 | 11,041 | 2,611 | 2,890 | 0.920 | 5.525 |
| | 3.82 | 4.31 | 26,674 | 7,164 | 6,184 | 0.859 | 5.371 |
| | 3.82 | 4.33 | 26,897 | 7,378 | 6,205 | 0.911 | 5.356 |
| Sp3+ComK | 4.04 | 4.30 | 17,133 | 3,808 | 4,393 | 1.210 | 5.328 |
| | 4.09 | 4.41 | 7,557 | 1,841 | 1,874 | 1.054 | 5.482 |
| Sp3+ComKΔ | 3.81 | 4.43 | 29,520 | 7,221 | 6,947 | 1.076 | 5.285 |
| | 3.90 | 4.43 | 29,343 | 7,739 | 6,301 | 1.062 | 5.076 |

**Extended Data Table 5. Parameters of fits of the FECS decays.** Local parameters specific to each independent experiment in the global fits are indicated with (1), (2), (3), (4) for experiments 1-4. Parameters without this annotation were global parameters.

| Construct | $a_1$ | $a_2$ | $\tau_1$ (ns) | $\tau_2$ (ns) | $a_3$ (1) | $a_3$ (2) | $a_3$ (3) | $a_3$ (4) |
|---|---|---|---|---|---|---|---|---|
| Construct: Box1-Sp-Box2 Fig. 4j | | | | | | | | |
| Box1 | 0.0172 | 0.00324 | 4.72 | 141.57 | 0.00345 | 0.00292 | - | - |
| Box1+ComK | 0.0163 | 0.00301 | 6.03 | 175.37 | 0.00306 | 0.00291 | - | - |
| Box1+ComKΔ | 0.0182 | 0.00365 | 4.95 | 143.58 | 0.00293 | 0.00281 | - | - |
| Sp | 0.0110 | 0.00164 | 4.16 | 154.00 | 0.00166 | 0.00218 | - | - |
| Sp+ComK | 0.0292 | 0.00401 | 5.63 | 171.51 | 0.00347 | 0.00342 | - | - |
| Sp+ComKΔ | 0.0200 | 0.00274 | 5.03 | 141.07 | 0.00263 | 0.00243 | - | - |
| Box2 | 0.0138 | 0.00231 | 4.18 | 135.56 | 0.00262 | 0.00269 | - | - |
| Box2+ComK | 0.0181 | 0.00254 | 5.90 | 159.72 | 0.00315 | 0.00250 | - | - |
| Box2+ComKΔ | 0.0165 | 0.00215 | 4.95 | 156.17 | 0.00250 | 0.00242 | - | - |
| Construct: Sp1-Sp2-Sp3-Box Fig. 5c | | | | | | | | |
| Box | 0.0080 | 0.00225 | 5.63 | 183.27 | 0.00287 | 0.00231 | 0.00323 | 0.00425 |
| Box+ComK | 0.0158 | 0.00421 | 6.00 | 145.90 | 0.00392 | 0.00354 | - | - |
| Box+ComKΔ | 0.0114 | 0.00216 | 6.52 | 220.76 | 0.00432 | 0.00399 | - | - |
| Sp3 | 0.0086 | 0.00178 | 4.52 | 139.86 | 0.00250 | 0.00242 | 0.00265 | 0.00283 |
| Sp3+ComK | 0.0185 | 0.00447 | 5.07 | 151.22 | 0.00381 | 0.00351 | - | - |
| Sp3+ComKΔ | 0.0141 | 0.00199 | 5.58 | 159.94 | 0.00342 | 0.00310 | - | - |
| Sp2 | 0.0097 | 0.00192 | 4.71 | 113.83 | 0.00252 | 0.00251 | 0.00313 | 0.00320 |
| Sp2+ComK | 0.0104 | 0.00258 | 5.6 | 141.07 | 0.00324 | 0.00369 | - | - |
| Sp2+ComKΔ | 0.0078 | 0.00204 | 6.14 | 227.57 | 0.00288 | 0.00310 | - | - |
| Sp1 | 0.0105 | 0.00317 | 5.67 | 208.09 | 0.00235 | 0.00150 | 0.00265 | 0.00265 |
| Sp1+ComK | 0.0129 | 0.00379 | 4.68 | 115.76 | 0.00320 | 0.00284 | - | - |
| Sp1+ComKΔ | 0.0108 | 0.00363 | 5.82 | 168.91 | 0.0271 | 0.00245 | - | - |
